# Haplotype and Population Structure Inference using Neural Networks in Whole-Genome Sequencing Data

**DOI:** 10.1101/2020.12.28.424587

**Authors:** Jonas Meisner, Anders Albrechtsen

## Abstract

Accurate inference of population structure is important in many studies of population genetics. Here we present, `HaploNet`, a method for performing dimensionality reduction and clustering of genetic data. The method is based on local clustering of phased haplotypes using neural networks from whole-genome sequencing or dense genotype data. By utilizing Gaussian mixtures in a variational autoencoder framework, we are able to learn a low-dimensional latent space in which we cluster haplotypes along the genome in a highly scalable manner. We demonstrate that we can use haplotype clusters in the latent space to infer global population structure utilizing haplotype information by exploiting the generative properties of our framework. Based on fitted neural networks and its latent haplotype clusters, we can perform principal component analysis and estimate ancestry proportions based on a maximum likelihood framework. Using sequencing data from simulations and closely related human populations, we demonstrate that our approach is better at distinguishing closely related populations than standard admixture and principal component analysis software. We further show that `HaploNet` is fast and highly scalable by applying it to genotype array data of the UK Biobank.

## Introduction

Understanding population structure is a cornerstone in population and evolutionary genetics as it provides insights into demographic events and evolutionary processes that have affected a population. The most common approaches for inferring population structure from genetic data are using principal component analysis (PCA) [41] and clustering algorithms such as `STRUCTURE` [42], or derivations thereof. PCA infers continuous axes of genetic variation that summarize the genetic relationship between samples while clustering algorithms assign samples to a fixed or variable number of ancestral sources while allowing for fractional membership. The inferred axes of PCA are very useful to account for population or cryptic structure in association studies or even to simply visualize the genetic data. A limitation for most PCA and clustering algorithms is that they assume all single-nucleotide polymorphisms (SNPs) to be independent, and they do therefore not benefit from the information of correlated sites or they may be biased thereof in their global estimates [55, 41]. A notable exception is `ChromoPainter` [32], which employs the Li and Stephens hidden markov model [33] for haplotype sampling in order to model and utilize correlations between SNPs, by letting samples be a mosaic of each other’s haplotypes. This has improved the fine-scale resolution and `ChromoPainter` has become state-of-the-art for inferring fine-scale population structure.

Gaussian mixture models and *k*-means are other commonly used methods for performing unsupervised clustering [48]. However, these methods suffer from the curse of dimensionality where relative distances between pairs of samples become almost indistinguishable in high-dimensional space [62]. A popular approach to overcome the curse of dimensionality is to perform dimensionality reduction, e.g. using PCA, and then perform clustering in the low-dimensional space that still captures most of the variation in the full dataset [15]. Recently, deep autoencoder methods have been very successful for large-scale datasets as they perform dimensionality reduction and clustering either sequentially or jointly to benefit from induced non-linearity and scalability of deep learning architectures using neural networks [60, 59]. Deep autoencoders have also been introduced in generative models, e.g. variational autoencoders, where the unknown data generating distribution is learnt by introducing latent random variables, such that new samples can be generated from this distribution [30, 45].

Most studies in population genetics utilizing neural networks for parameter inference have mainly focused on supervised learning through simulations from demographic models [50, 49, 17, 18, 10]. Here, an overall demography is assumed, based on previous literature, and a lot of different datasets are simulated using small variations in model parameters, e.g. selection coefficient or recombination rate, with evolutionary simulators (e.g. `msprime` [26] or `SLiM` [20]). The studies usually convert a simulated haplotype matrix into a downscaled fixed sized image with rows and/or columns sorted based on some distance measure. The network is then trained on the simulated datasets to learn the specified model parameters with feature boundaries in convolutional layers, and in the end, the model is tested on a real dataset. However, recently more studies have instead focused on deep generative models for data-driven inference or simulation using unsupervised learning approaches which will also be suitable for the growing number of unlabelled large-scale genetic datasets [39, 61, 5, 58, 3]. There has also been a recent interest in the application of the non-linear dimensionality reduction method UMAP in population genetics [13].

We here present `HaploNet`, a method for inferring haplotype clusters and population structure using neural networks in an unsupervised approach for phased haplotypes of whole-genome sequencing (WGS) or genotype data. We utilize a variational autoencoder (VAE) framework to learn mappings to and from a low-dimensional latent space in which we will perform indirect clustering of haplotypes with a Gaussian mixture prior. Therefore, we do not have to rely on simulated training data from demographic models with a lot of user-specified parameters but are able to construct a fully data-driven inference framework in which we can infer fine-scale population structure. We locally cluster haplotypes and exploit the generative nature of the variational autoencoder to perform PCA and to build a global clustering model similar to `NGSadmix` [53], which we fit using an accelerated expectation-maximization (EM) algorithm to estimate ancestry proportions as well as frequencies of our neural network inferred haplotype clusters.

## Results

### Simulation scenarios

We performed a simulation study of five populations inspired by the simple demography in [32] and applied `HaploNet` to four different scenarios. The demography used for simulation is displayed in Figure 1A. We simulated three different scenarios of various population split times with a constant population size of 10,000 as well as another scenario with a constant population size of 50,000, to counteract the increased genetic drift in an increase of population split times. In all scenarios, we simulated 500 diploid individuals with 100 from each of the five populations. We compare the estimated ancestry proportions from our `HaploNet` clustering algorithm against results from `ADMIXTURE` based on signal-to-noise ratio measures (Table S1) and their ability to separate ancestry sources. We further compare the inferred population structure from our PCA approach to the PCA from `ChromoPainter` and standard PCA using `PLINK` based on signal-to-noise ratio measures of the inferred principal components (Table S2). The results of the ancestry estimation of all simulation scenarios are shown in Figure S2.

**Figure 1:**
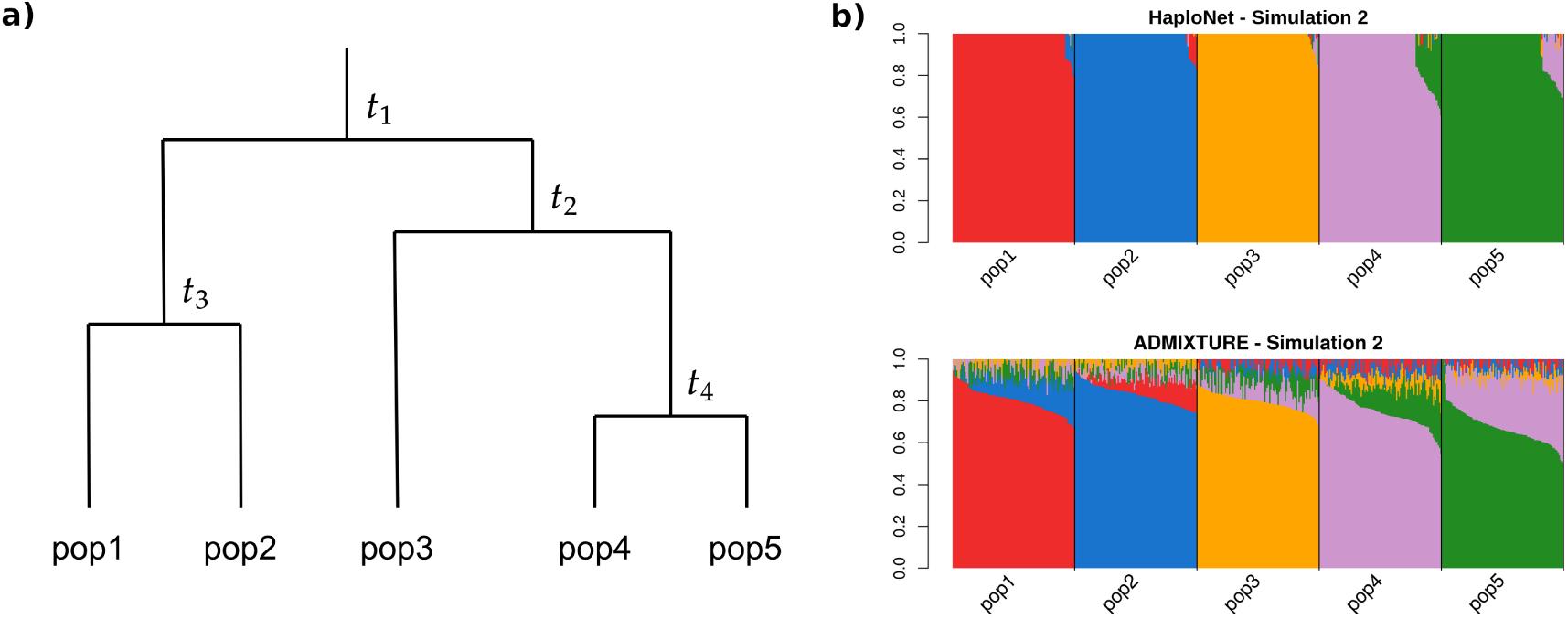
***a)*** Overview of simulation configuration of four splits into five populations with equal population sizes at all times. The time of population splits are designated *t*_1_, *t*_2_, *t*_3_ and *t*_4_, measured in generations. ***b)*** Estimated ancestry proportions in one of the four simulation scenarios (Simulation 2) with *t*_1_ = 200, *t*_2_ = 120, *t*_3_ = 80 and *t*_4_ = 40 using `HaploNet` (top) and `ADMIXTURE` (bottom).

Simulation 1 is the hardest scenario where the split times measured in generations between populations are very recent, *t*_1_ = 100, *t*_2_ = 60, *t*_3_ = 40 and *t*_4_ = 20 with a constant population size of 10,000. Here, we see `HaploNet` capturing more structure and are able to better separate the five populations than `ADMIXTURE` as also seen in the signal-to-noise ratio measures. As expected due to the recent split between pop4 and pop5 have not had time to become two distinct homogeneous populations and we are not able to perfectly separate the populations into homogeneous clusters. For the PCAs, visualized in Figure S3, we see that all three methods are able to split the five populations with only small overlap between few individuals of pop4 and pop5 on PC4.

In Simulation 2 and 3, the split times are respectively 2 and 10 times longer than in Simulation 1 and in these easier scenarios, the `ADMIXTURE` proportion estimates from `ADMIXTURE` are still very noisy whereas `HaploNet` has better separation of the populations (Figure 1B and S2). All methods are able to infer distinct clusters of the populations using PCA (Figure S4 and S5) with `ChromoPainter` having the best signal-to-noise ratio for scenario 2 and a similar performance to `HaploNet` in scenario 3 (Table S2).

Simulation 4 has the same split times as Simulation 3 (*t*_1_ = 1000, *t*_2_ = 600, *t*_3_ = 400 and *t*_4_ = 200) but with a with a constant population size of 50,000, which lower the effect of genetic drift that makes the populations less distinct, and thus makes the scenario harder than Simulation 3. We observe that `HaploNet` is still capable of perfectly separating the ancestry sources while `ADMIXTURE` has noisy estimates for all individuals. We note that `ChromoPainter` was prematurely terminated due to memory error for 13.4 million SNPs by exceeding the available memory on the test machine (128 GB) and the expected runtime would have been approximately 1.5 month. `HaploNet` and `PLINK` are able to perfectly split and cluster the populations in PC space as visualized in Figure S6. In all scenarios, we performed the `ADMIXTURE` analyses and the `PLINK` PCA with and without LD pruning. For all analyses the method performed slightly better with LD-pruned data (Table S1 and S2).

### 1000 Genomes Project

We applied `HaploNet` to the entire 1000 Genomes Project and separately to each its five super populations (AFR, AMR, EAS, EUR, SAS). Each super population had between 347 and 661 individuals and 3.2-8.4 common (MAF *>* 5%) SNPs (Table 2). We compare the estimated individual ancestry proportions from our clustering algorithm with results from `ADMIXTURE` based on signalto-noise ratio measures (Table S3), and we compare the inferred population structure from our PCA approach with the PCA from `ChromoPainter` and standard PCA using `PLINK` as well based on signal-to-noise ratio measures of the inferred principal components (Table S4) for each of the super populations. Additionally, we compare the runtimes of `HaploNet` and `ChromoPainter` for chromosome 2 in each of the five super populations using both GPU and CPU for training `HaploNet`, which is summarized in Table 1. Overall illustrations of ancestry estimations for the full dataset (*K* = 15) and for each super population are depicted in Figure 2 and 3, respectively. The application of `HaploNet` on the full dataset has been performed to validate the fine-scale structure in the super population applications. Another visualization of ancestry estimations in the full dataset is shown in Figure S23 for *K* = 14 that was the highest *K* for which `ADMIXTURE` has reached convergence.

**Table 1:** Runtimes for chromosome 2 in the five super populations of the 1000 Genomes Project using `HaploNet` and `ChromoPainter`. `HaploNet` has both been trained on a GPU and a CPU on a machine with 24 threads. In all scenarios, models were trained with a fixed window size of 1024 SNPs, a batch-size of 128 for 200 epochs.

|  | **HaploNet (GPU)** | **HaploNet (CPU)** | **ChromoPainter** |
| --- | --- | --- | --- |
| **AFR** | 1.8 hours | 3.9 hours | 95.7 hours |
| **AMR** | 0.7 hours | 1.6 hours | 21.8 hours |
| **EAS** | 0.8 hours | 1.9 hours | 41.7 hours |
| **EUR** | 0.9 hours | 2.1 hours | 45.6 hours |
| **SAS** | 0.9 hours | 2.1 hours | 41.9 hours |

**Table 2:** General dataset information of the super populations in the 1000 Genomes Project and the number of windows and their mean size used by `HaploNet`.

| Population | #Individuals | #SNPs | #Windows | Mean window size (in Mb) |
| --- | --- | --- | --- | --- |
| AFR | 661 | 8.4 million | 8239 | 0.34 |
| AMR | 347 | 3.2 million | 6071 | 0.46 |
| EAS | 504 | 5.6 million | 5450 | 0.51 |
| EUR | 503 | 6.0 million | 5904 | 0.47 |
| SAS | 489 | 6.2 million | 6054 | 0.46 |
| Full | 2504 | 6.9 million | 6705 | 0.41 |

**Figure 2:**
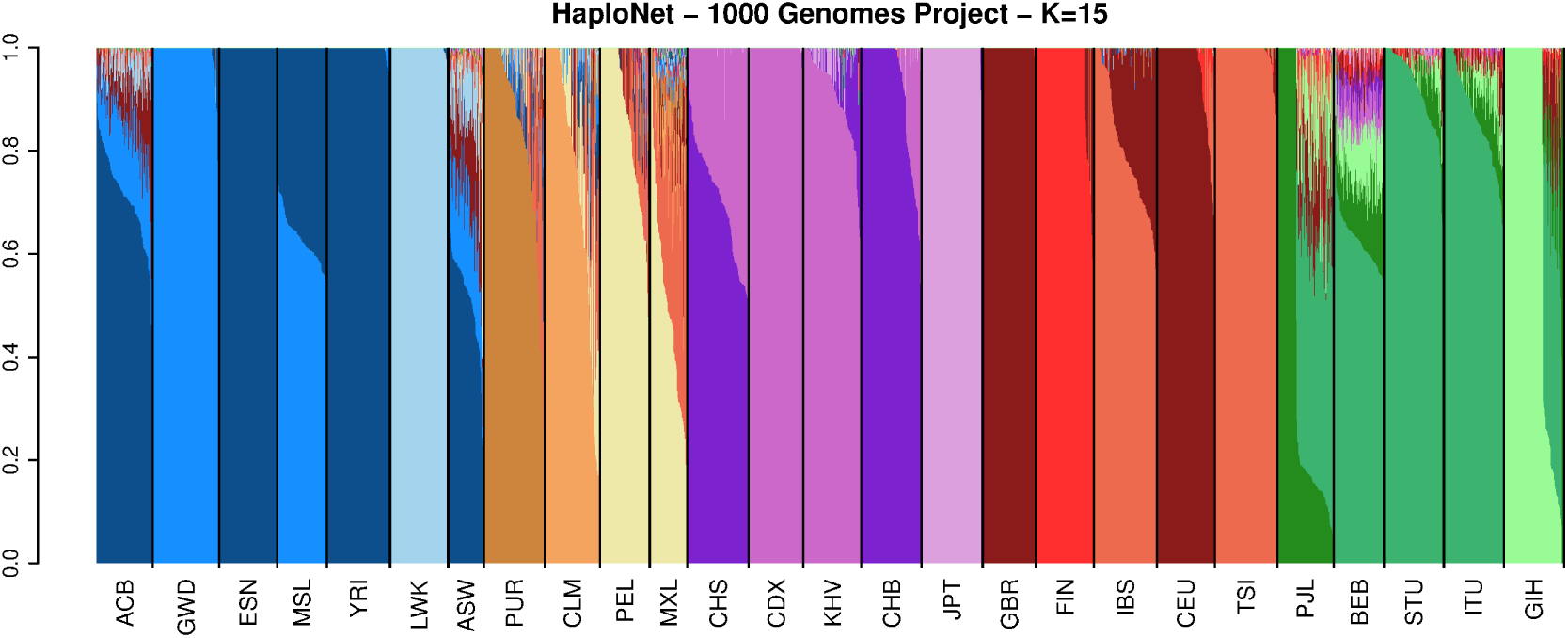
Estimated ancestry proportions in the full 1000 Genomes Project using `HaploNet` for *K* = 15. `ADMIXTURE` was not able to converge to a solution in 100 runs for this scenario.

**Figure 3:**
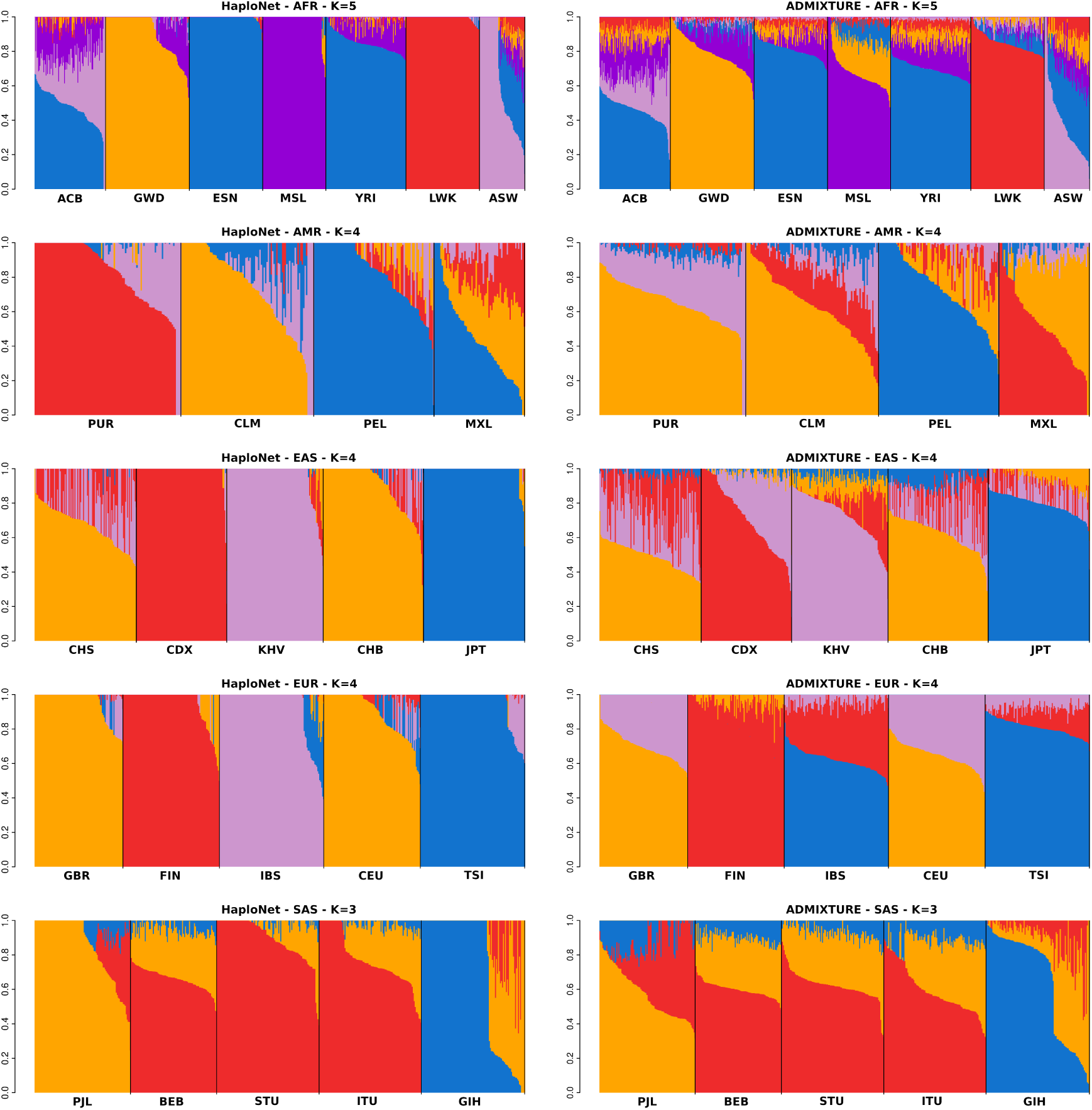
Estimated ancestry proportions in the super populations of the 1000 Genomes Project using `HaploNet` (left column) and `ADMIXTURE` (right column) for AFR, AMR, EAS, EUR, SAS, respectively.

The African (AFR) super population includes 661 individuals from seven populations with two populations also having European ancestry (African Caribbean, ACB, and African American, ASW). The inferred population structure for different *K* and PCs are shown in Figure S7 and S8. The benefit of using haplotype information is immediately clear, as the estimated ancestry proportions from `HaploNet` are much better at separating the populations as also seen by the signal-to-noise ratio measure in comparison to `ADMIXTURE`. This is clearly visible for *K* = 5 (Figure 3) where the MSL population is represented by its own component and its ancestry is also seen in the GWD and YRI populations, which makes sense from a geographical viewpoint. For PCA (Figure S8), the beneficial effect of using the haplotype-based methods is not apparent as the large scale structure from the European ancestry within ACB and ASW makes it hard to distinguish fine-scale structure within the African ancestry. This European ancestry is correctly inferred in the full dataset with *K* = 15 (Figure 2) where both ASW and ACB have substantial north European ancestry (GBR and CEU).

The results of the American (AMR) super population containing 347 individuals are visualized in Figure S9 and S10. These population consists of many individuals with both European and Native American ancestry. Interestingly, we observe that `HaploNet` and `ADMIXTURE` clusters the individuals differently, where `HaploNet` splits the PUR and CLM populations that could correspond to two different European ancestry sources while `ADMIXTURE` splits the Native American ancestry within the PEL and MXL populations, for *K* = 3, 4. In *K* = 4, we see that `HaploNet` captures an African ancestry signal in PUR and to a lesser degree in CLM. This is consistent with previous studies [19] and validated from running on the full dataset (Figure 2). In the full data, we can also see that the European ancestry in the Mexican population (MXL) is mainly from southern European (TSI and IBS) while the PUR and CLM European ancestry gets its own component. In the PCA plots (Figure S10), we see that `HaploNet` and `ChromoPainter` are clustering the populations slightly better than standard PCA by making them more separable and separating the signal captured by PC3 and PC4.

We next analyzed the 504 individuals in the East Asian (EAS) super population (Figure 3, Figure S11 and S12). From the estimated ancestry proportions `HaploNet` perform much better than `ADMIXTURE` and is able to cleanly separate the Vietnamese ancestry (KHV) from the ancestry signal of the Dai people (CDX) for *K* = 4. We see a very similar pattern on the PCA plots, where the haplotype based methods are able to separate the two populations as well, while the standard PCA approach can not.

The results of the European (EUR) super population are visualized in Figure S13 and S14 for ancestry proportions and PCA, respectively. For *K* = 4 (Figure 3), we observe that `HaploNet` is able to distinguish between the two Southern European populations (IBS and TSI) which is not the case for `ADMIXTURE`. We also see from the signal-to-noise ratio measures that `HaploNet` is much better at separating the five populations. A similar pattern is observed in the population structure inferred using PCA for `HaploNet` and `ChromoPainter` where they are able to separate the Southern European populations on PC3 as also verified with their signal-to-noise ratio measures.

Finally, the results South Asian (SAS) super population are visualized in Figure S15 and S16. We see that the complex population structure of the SAS super population makes it hard for the haplotype based methods to separate the populations on a finer-scale than the standard approaches.

The presented analyses were run on computer clusters and in order to benchmark the runtimes, we timed the analyses of chromosome 2 on a single machine with both CPU and GPU. `HaploNet` was about twice as fast using the GPU compared to CPU and `ChromoPainter` was 10-30 times slower (Table 1), and we were not able to run it on the full 1000 Genomes Project data.

### Robustness

To test the robustness of `HaploNet` in terms of hyperparameters, model and data type, we applied it to the European super population under various scenarios and compared the performance to the results above. The performances are summarized in Table S5.

We varied window sizes and evaluated its performance on inferring population structure based on both ancestry proportions and PCA. The results are displayed in Figure S17 and S18. `HaploNet` estimates very similar ancestry proportions for each of the window sizes where we are able to separate the Southern European populations. However, we see a decreasing degree of resolution in the ancestry proportions as the window size becomes larger. For the PCA plots, we observe that all windows sizes are able to separate the Southern European populations and have similar performances in terms of signal-to-noise ratio measures.

We investigated the effectiveness of the Gaussian mixture prior model by only having a model with a categorical latent variable and a linear decoder. The model would then perform amortized haplotype clustering with the decoding process directly modelling haplotype cluster frequencies. The results are shown in Figure S19, where we see a decrease in the model’s sensitivity to separate the Southern European populations in comparison to our full model.

We also verified the model’s dependency on LD structure to infer fine-scale population structure. Here, we permuted SNP positions of all chromosomes to distort and remove the LD structure in the data. The results are visualized in Figure S20, where the model is not able to infer any meaningful population structure further than on PC1.

Lastly, we analyzed only the SNPs found on the high density genotype chip of the Omni platform [1], to test how well `HaploNet` would perform on SNP array data. After filtering, the dataset contained of 1.3 million SNPs and we have used a window size of 256 in `HaploNet`. The results are visualized in Figure S21 and S22, where `HaploNet` has similar performance in comparison to when it is used on the full dataset and is also fully able to split the two Southern European populations which was not possible for `ADMIXTURE` with the full dataset.

### UK Biobank

We further applied `HaploNet` to SNP array data of the UK Biobank dataset and infer population structure on a subset of 276,732 unrelated with self-reported ethnicity as ‘White-British’ with a total of 567,499 SNPs after quality control and filtering. It took 7.3 hours to train `HaploNet` on the chromosome 2 with a window size of 256, a batch-size of 8196 for 100 epochs. The results are displayed in Figure 4 and S25 for estimating ancestry proportions and inferring population structure using PCA, respectively. For *K* = 3, `HaploNet` infers three clear ancestral components that reflect English, Scottish and Welsh sources, while for *K* = 4, we infer an additional ancestral component capturing a signal in North West England. We have visualized the distributions of ancestry proportions stratified by country of birth for *K* = 4 in Figure S24. For the population structure inferred using our PCA method, we capture similar structure in the top PCs, where we additionally see a component capturing the variation between North and South Wales. On PC6, we capture structure that does not reflect population structure but instead captures variation that could be due to SNP ascertainment or strong LD structure as previously observed for the UK Biobank [43]. The signal is caused by a single genomic region as shown by the SNP loadings in Figure S26. Similar cluster patterns in the North West England have previously been observed before [47] in the ‘White-British’ subset of the UK Biobank.

**Figure 4:**
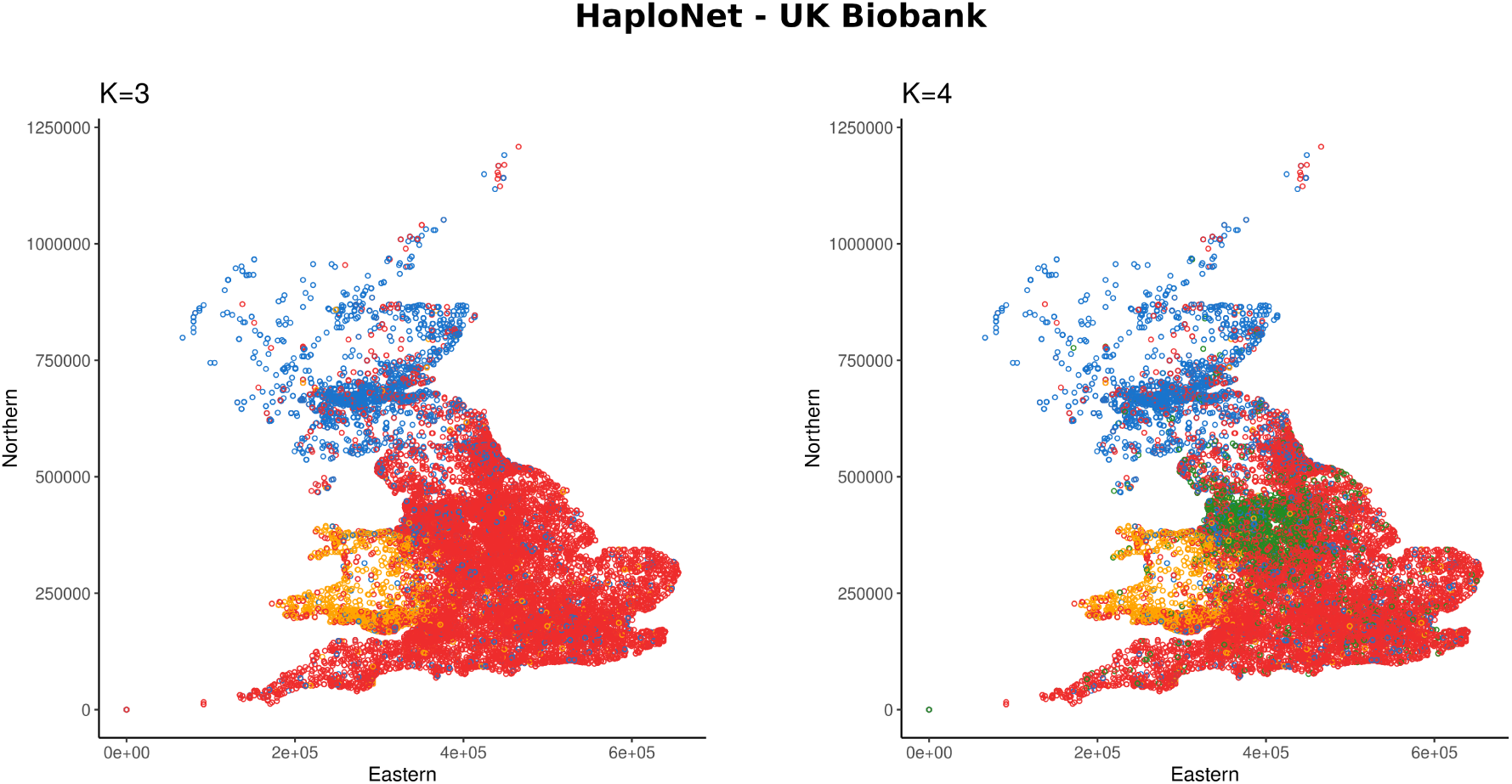
Estimated ancestry proportions in the subset of unrelated self-identified ‘White-British’ of the UK Biobank using `HaploNet` for *K* = 3 and *K* = 4, respectively. Individuals are plotted by their birthplace coordinates and colored by their highest associated ancestry component.

## Discussion

We have presented our new framework, `HaploNet`, which performs dimensionality reduction and clusters haplotypes using neural networks. We explored its capability to infer population structure based on local haplotype clusterings, which are performed in windows along the genome, as well as its generative properties for estimating neural network likelihoods. We show the benefits of merging machine learning with traditional statistical frameworks, as we developed a novel method for estimating ancestry proportions from neural networks in a likelihood model. We tested `HaploNet` in simulation scenarios and data from the 1000 Genomes Project and compared its results to commonly used software for inferring population structure based on ancestry proportions or PCA. We demonstrated that `HaploNet` is capable of utilizing haplotype information for inferring fine-scale population structure that outperforms `ADMIXTURE` with respect to signal-to-noise ratio. On real datasets, `HaploNet` infers similar population structure to `ChromoPainter` using PCA, while being much faster. However, in several simulations `ChromoPainter` separated the populations better than `HaploNet`. It was not computationally feasible to run `ChromoPainter` for larger data sets both in terms of speed and memory usage. We further showed the scalability of `HaploNet` by applying it to genotype data of the UK Biobank with hundreds of thousands of individuals, which is not possible with `ADMIXTURE` or `ChromoPainter`, and we are able to infer four ancestry sources for individuals born within the United Kingdom.

We have also reported the performances of `ADMIXTURE` and `PLINK` on a LD pruned version of all datasets for ancestry estimation and PCA, respectively. In all cases, we observed similar or slightly better performance in comparison to using the full non-pruned dataset. However, the consequence of LD pruning in a heterogeneous dataset with population structure besides losing information is not fully understood because the population structure will increase the LD between SNPs. LD pruning can greatly decrease *F*_*ST*_ values between populations [34] and thus affects the distribution of eigenvalues since they are proportional to *F*_*ST*_ [37]. Therefore, we have measured the accuracy of PCA (`PLINK`) and `ADMIXTURE` with and without pruning for each of the super populations. However for computational reasons, we have only used a LD pruned version when analyzing the full 1000 Genomes Project data with `ADMIXTURE`. Another factor that can affect measures of population structure is minimum minor allele frequency cutoffs. We have chosen the standard 5% cutoff for most analyses including `ADMIXTURE` and PCA in `PLINK`. Rare alleles can reduce the performance of these methods [35] and are harder to phase which can be a problem for `HaploNet` and `ChromoPainter`. On the other hand they can also be informative about more recent population structure. However, we have not explored the effect of including the more rare alleles in this study.

The number of clusters, *C*, is usually a non-trivial hyperparameter to set in a Gaussian mixture model or in a genetic clustering setting. Multiple runs of varying *C* are usually performed and evaluated based on some criteria. In our study, we saw that `HaploNet` seemed capable of inferring the optimal number of haplotype clusters to use by setting *C* to a high fixed number (*C* = 32 in all analyses). At lower *C*, e.g. 20, we observed a slightly worse performance in the analysis of the African super population of the 1000 Genomes Project but similar performance for the other super populations (not shown). It therefore seems that choosing a high value for *C* does not decrease the performance for super populations such as the European which has less genetic diversity. In those cases, `HaploNet` will only use a subset of the possible haplotype clusters to model the haplotype encodings, which has also been observed in a different application of the GMVAE model [9].

A limitation of our model and other deep learning models, is that usually only relatively small genetic datasets are available to researchers, which introduces problems with training convergence and overfitting. To combat these issues, we kept the number of parameters low and used an autoencoder architecture that naturally regularizes its reconstruction performance. Another advantage of having a small model configuration is observed for the low training times on both GPU and CPU setups that broadens the application opportunities of `HaploNet`. However, the difference between the GPU and CPU runtimes will be larger when running chromosomes in parallel. For all analyses in this study including the UK Biobank data, we have been using the entire data for training our neural networks to model all available haplotypes, whereas with larger labelled datasets, one could have a separate training data. We have also limited ourselves to using fixed window lengths across chromosomes for the matter of simplicity and ease of use, where we instead could have included external information from genetic maps to define windows of variable size. We show that `HaploNet` is somewhat robust to changes in window size when inferring global population structure, either using PCA or ancestry proportions, however for fixed-size windows, there is a trade-off in resolution and training time that is subject for future research. We further show that we are able to capture fine-scale structure when only evaluating variable sites available on a common genotype chip that allows for broader applications of our method.

Our model serves as a proof-of-concept and an exploration for how non-linear neural networks and specialized architectures can be utilized to learn haplotype clusters and encodings across a full genome in a very scalable procedure. We hypothesize that as the number of large-scale genetic datasets are growing, we will see the increasing importance of deep learning in population genetics, as deeper models can be trained and more bespoke architectures will be developed. As shown in our study, we can even use learnt mappings together with standard statistical frameworks to further improve our understanding of genetic variation. Future developments of our framework is to use the haplotype clusters in sequential models, to e.g. infer local ancestry and IBD tracts in HMMs, as well as investigate its potential integration into imputation based on haplotype reconstruction and association studies.

## Methods

The method is based on phased haplotype data from diallelic markers. We define the data matrix **X** as a 2*N* × *M* haplotype matrix for a given chromosome, where *N* is the number of individuals and *M* is the number of SNPs along the chromosome. The entries of the matrix are encoded as either 0 or 1, referring to the major and minor allele, respectively.

For each chromosome, we divide the sites into *W* windows of a fixed length of *L* SNPs, which we assume is much smaller than *M*. The windows are non-overlapping and we will further assume that the parameters estimated in a window are independent from parameters estimated in adjacent windows. The length of the genomic windows can also be defined by a recombination map, however we have kept it fixed for the sake of generalizability and ease of application in this study. For each defined window along a chromosome, we independently train neural networks in a variational autoencoder framework to learn haplotype clusters and encodings using a Gaussian mixture prior. We define haplotype clusters as a collection of haplotypes that cluster together based on similarities in their encodings. In the model, they are identified by their latent state (*y*) whose mean structure predicts a distribution of similar haplotypes. We are then able to calculate a likelihood for an observed haplotype given an inferred haplotype cluster which resembles calculating genotype likelihoods in WGS data from the trained networks.

### Variational autoencoder

An autoencoder is a state-of-the-art approach for performing dimensionality reduction by learning a mapping of the space of the input data, *χ*, to a low-dimensional space 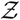 and a mapping back to the input data [46, 4]. More formally, we can describe the two mapping functions as 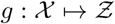 and 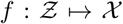, which are commonly called the encoder and the decoder, respectively. Both the encoder and the decoder are parameterized by (deep) neural networks to learn the mappings, as multilayer feed-forward neural networks are universal function approximators [22].

A probabilistic variant of this neural network architecture is introduced in the variational autoencoder (VAE), where the unknown generating process of the input data is modelled by introducing latent variables and the joint probability is approximated through variational inference. As an optimization method, variational inference is often used to approximate the posterior distribution of a set of latent variables **z**, *p*(**z** | **x**), by fitting a function that describes a chosen family of distributions, *q*_*ϕ*_(**z** | **x**). Thus, variational inference turns it into an optimization problem, where the objective is to maximize the evidence lower bound (ELBO) of the marginal log-likelihood of the data, *p*_*θ*_(**x**), iteratively. In contrast, Monte Carlo Markov Chain methods approximate the joint probability of the data and the latent variables by sampling from the posterior distribution [7]. The function approximating the posterior distribution is parameterized with variational parameters *ϕ*. Kingma and Welling introduced the Stochastic Gradient Variational Bayes (SGVB) estimator [30] of the ELBO for approximate posterior inference in a VAE framework (as well as [45]), where a set of parameters, (*θ, ϕ*), are optimized with amortized inference using mappings parameterized by neural networks. Here, the marginal log-likelihood of the data, *p*_*θ*_(**x**), is parameterized with parameters *θ*. This amortization means that the number of parameters does not depend on sample size as in traditional variational inference but depends on the network size [52]. The VAE can be seen as an autoencoder with its latent space being regularized by a chosen prior distribution to make the inferred latent space more interpretable and to preserve global structure. A standard Gaussian prior is the most common choice, however, it is often too simple and a lot of effort has been made to make the approximate posterior richer with normalizing flows and additional stochastic layers [44, 29, 54].

In our proposed method, `HaploNet`, we construct a generative model and use a variational autoencoder framework to learn low-dimensional encodings of haplotypes in windows along the genome. However, we also introduce an additional categorical variable *y* to represent haplotype clusters as mixture components such that we assume a Gaussian mixture prior in the generative model. In this way, we are able to jointly perform dimensionality reduction and cluster haplotypes in a highly scalable approach for a given genomic window. In the following model descriptions, we will follow the mathematical notation used in the machine learning literature and Kingma and Welling (2013) [30], where *p*_*θ*_ and *q*_*ϕ*_ are probability functions that define the decoder and encoder part, respectively. *θ* and *ϕ* are the parameters (biases and weights) in the neural networks. We have provided a visualization of our overall network architecture, including descriptions of the major substructures, in Figure 5. We define the following latent variable model for a single haplotype in a single genomic window with data **x** *∈ {*0, 1*}*^*L*^, Gaussian latent variables **z** *∈* ℝ^*D*^ and categorical latent variable *y ∈ {*0, 1*}*^*C*^ (one-hot encoded) such that **x** is conditionally independent of *y*:

**Figure 5:**
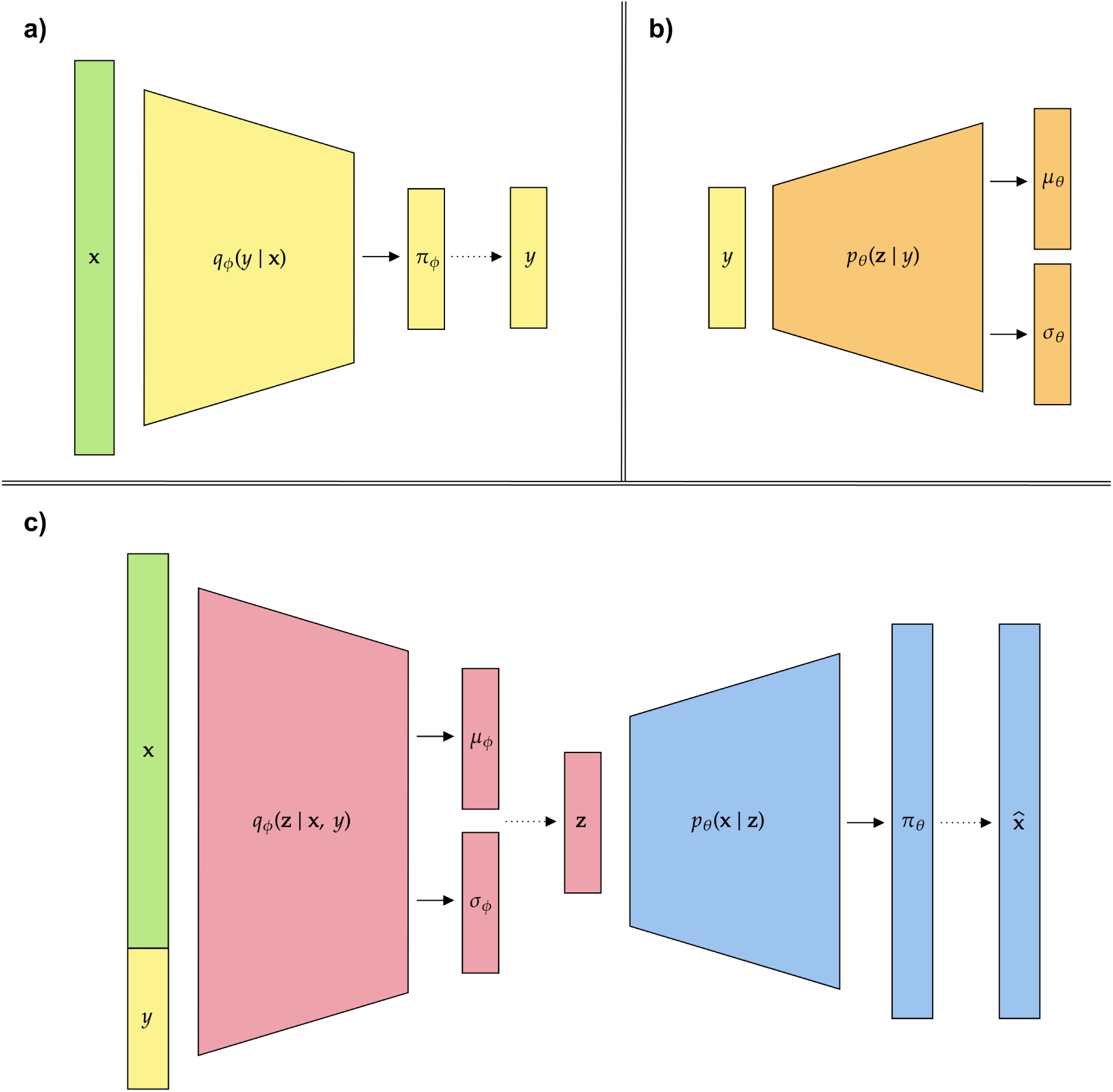
The neural network architecture of `HaploNet` split into three major substructures. Here the solid lines represent the estimation of distribution parameters, while the dashed lines represent sampling of latent variables. ***a)*** Displays the neural network parameterizing the distribution *q*_*ϕ*_(*y* | **x**), for sampling the haplotype cluster, ***b)*** the network parameterizing the regularizing distribution of the sampled encoding, *p*_*θ*_(**z** | *y*), and ***c)*** shows the network parameterizing the distribution *q*_*ϕ*_(**z** |**x**, *y*), for sampling the haplotype encoding, as well as the network decoding the sampled encoding to reconstruct our input. Note that the colors of the network blocks are coherent across substructures such that the sampled *y* in ***a)*** is used in both ***b)*** and ***c)***.

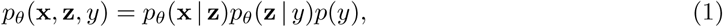

with generative processes defined as:

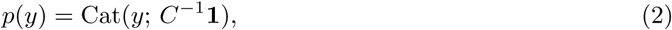

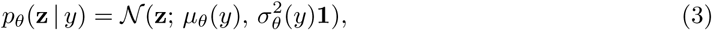

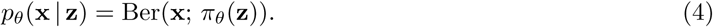

Here **z** is a *D*-dimensional vector representing the latent haplotype encoding and *C* is the number of haplotype clusters, while 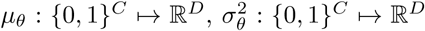 are mapping functions parameterized by neural networks with network parameters *θ*. In this case, Ber(**x**; *π*_*θ*_(**z**)) is a vectorized notation of Bernoulli distributions and each of the *L* sites will have an independent probability mass function. We assume that the covariance matrix of the multivariate Gaussian distribution is a diagonal matrix which will promote disentangled factors. We assume the following inference (encoder) model that constitutes the approximate posterior distribution:

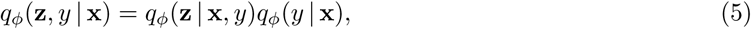

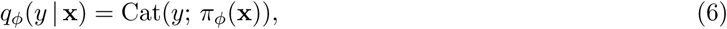

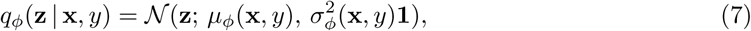

where 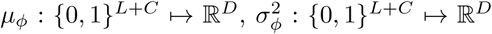 and *π*_*ϕ*_ : {0, 1} ^*L*^ *→* [0, 1]^*C*^ again are mapping functions parameterized by neural networks with network parameters *ϕ*. Therefore, the marginal posterior distribution and marginal approximate posterior distribution of **z** will both be a mixture of Gaussians. Thus, *q*_*ϕ*_(**z**, *y* | **x**) and *p*_*θ*_(**x** | **z**) will constitute the probabilistic encoder and decoder, respectively, in comparison to the deterministic encoder and decoder of the standard autoencoder.

From the marginal log-likelihood of the data, we derive the following ELBO [8] of our variational autoencoder model for haplotype *i*,

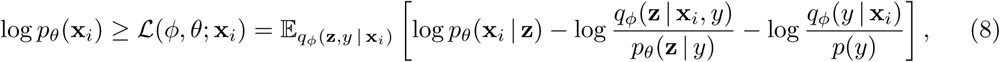

where the marginal log-likelihood of the full data in a window is given by:

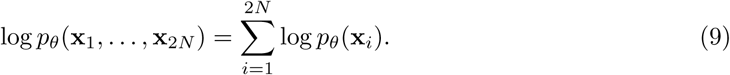

The full derivation of this ELBO is described in the supplementary material as well as reparameterization tricks that are needed to approximate and optimize it through Monte Carlo samples of the latent variables. We immediately see that the first term in Equation 8 describes the reconstruction error of mapping from the latent space back to the input space as in an autoencoder framework. The next two terms act as regularization on the learnt latent spaces, where the second term encourages the variational Gaussian distributions to be close to the estimated prior distributions, while the last term encourages anti-clustering behaviour to prevent all haplotypes to cluster in one component. This is a modification of the unsupervised loss of the M2 model [28], as described by Rui Shu [51], where information of the haplotype cluster is also propagated through *p*_*θ*_(**z** | *y*). However, we further approximate the categorical latent variable with samples from a Gumbel-Softmax distribution [24, 36] instead of the categorical distribution. The Gumbel-Softmax distribution is a continuous approximation to the categorical distribution that can be easily reparameterized for differentiable sampling and gradient estimations (Figure S1). In this way, we can avoid an expensive computational step of having to marginalize over the categorical latent variable in the SGVB estimator of the ELBO as is done in the original model. A lot of different interpretations and implementations of the Gaussian Mixture Variational Autoencoder (GMVAE) have been proposed [14, 12, 9, 25], and a similar architecture to ours has been implemented [16].

### Neural network likelihoods

We can exploit and utilize the generative nature of our GMVAE model based on the parameters of the trained neural networks for a window to construct likelihoods from the mean latent encodings of the estimated haplotype clusters through reconstruction. We define 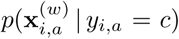 with *y*_*i,a*_ being one-hot encoded, as the neural network (NN) likelihood that the data is generated from the *c*-th haplotype cluster, for the *a*-th haplotype of individual *i* in window *w*. The NN likelihoods are calculated as follows using the probability mass function of the Bernoulli distribution and the properties, 𝔼 [*p*_*θ*_(**z** | *y*)] = *µ*_*θ*_(*y*) and 𝔼 [*p*_*θ*_(**x** | *µ*_*θ*_(*y*))] = *π*_*θ*_(*µ*_*θ*_(*y*)):

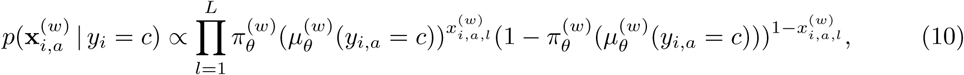

for *c* = 1, …, *C* (one-hot encoded) and 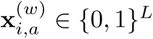 is the data of the *a*-th haplotype of individual *i* in window *w* with *L* being the number of SNPs. Here 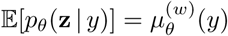 describes the learnt mean latent encoding of a given haplotype cluster, *y*, in the generative model and 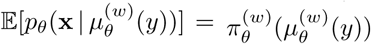 is the learnt mean reconstruction of the mean latent encoding.

### Ancestry proportions and haplotype cluster frequencies

A widely used approach for inferring population structure is estimating ancestry proportions. We propose a model for estimating ancestry proportions and haplotype cluster frequencies assuming *K* ancestral components based on the model introduced in `NGSadmix` [53], as an extension to the `ADMIXTURE` model [2], where instead of latent states of unobserved genotypes, we have latent states of haplotype clusters. We can then construct the following likelihood model of the data **X** using the above defined NN likelihoods given the genome-wide ancestry proportions **Q** and window-based ancestral haplotype frequencies **F**:

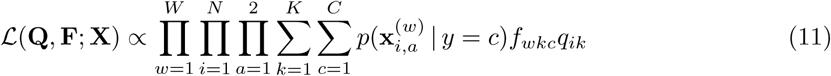

with *k* describing the ancestral state, for **Q** *∈* [0, 1]^*N×K*^ with constraint 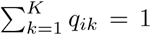 with constraint 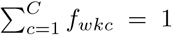. Here *W* is the total number of windows across chromosomes, *N* is the number of individuals and *C* is the number of haplotype clusters. Maximum likelihood estimates of F and Q are obtained using an expectation-maximization (EM) algorithm. The full description of the EM algorithm is detailed in the supplementary material. We use the S3 scheme of the SQUAREM methods [57] for accelerating our EM implementation, such that one large step is taken in parameter space based on the linear combination of two normal steps.

### Inference of population structure using PCA

We can also use the NN likelihoods to infer population structure using PCA, however, the approach is not as straightforward as for the model-based ancestry estimation. We use a similar approach as in microsatellite studies where we assume that all clusters are an independent marker, and we simply sum the cluster counts for each individual in each window by taking the most probable cluster for each haplotype to construct a *N × W × C* tensor, **Y**.

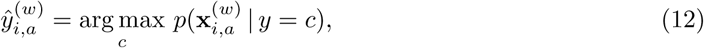

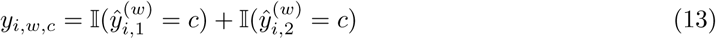

We can now treat the task as a standard PCA approach in population genetics based on a binomial model [41] such that the pairwise covariance is estimated as follows for individual *i* and *j*:

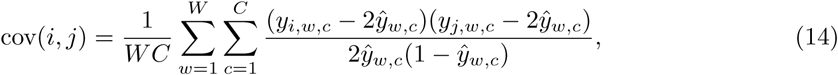

where 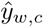 is the frequency of the *c*-th haplotype cluster in window *w*. We can finally perform eigendecomposition on the covariance matrix to extract principal components.

### Implementation

We have implemented `HaploNet` as a Python program using the `PyTorch` library (v.1.10) [40], and it is freely available on https://github.com/rosemeis/HaploNet. We have used the `NumPy` [21] and `scikit-allel` [38] libraries for preprocessing the data from Variant Call Format (VCF) into data structures to be used in `HaploNet`. The EM algorithm for estimating ancestry proportions and the algorithm for performing PCA have been implemented in Cython [6] for speed and parallelism.

An overall detailed description of the network architectures used in different scenarios can be found in the supplementary material. We have used fully-connected layers throughout the network, and for all inner layers in our neural networks, we are using rectified linear unit (ReLU(*x*) = max(0, *x*)) activations to induce non-linearity into the networks, followed by batch normalization [23], while all outer layers are modelled with linear activations. The usage of linear activations means that the networks are estimating the logits of the probabilities instead the probabilities directly in *π*_*θ*_(**z**) and *π*_*ϕ*_(**x**) for computational stability, as well as for 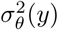 and 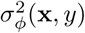 that represent log *σ*^2^ in inner computations.

We are training our networks with the Adam optimizer [27] using default parameters with a learning rate of 1.0 × 10^*−*3^, *β*_1_ = 0.9 and *β*_2_ = 0.999. The batch-sizes and number of epochs are detailed in the supplementary material for each of the different datasets. We have used a fixed temperature in the sampling from the Gumbel-Softmax distribution of *τ* = 0.1 to approximate and encourage a categorical sampling, and we use one Monte Carlo sample of the latent variables to approximate the expectation in equation 8.

### Simulations

We have simulated five populations from a simple demography with four population splits (*t*_1_, *t*_2_, *t*_3_, *t*_4_) using `msprime` [26] (Figure 1A). We have simulated 500 diploid individuals in four different scenarios with various changes to the chosen population split times and the population sizes. A uniform recombination rate of 1.0 *×* 10^*−*8^ has been assumed in all simulations as well as a mutation rate of 2.36 *×* 10^*−*8^ [56] for a sequence of 1.0 *×* 10^9^ bases.

Simulation 1 has the following population split times: *t*_1_ = 100, *t*_2_ = 60, *t*_3_ = 40 and *t*_4_ = 20 with an assumed constant population size of 10,000 at all points. After filtering for minor allele frequency threshold of 0.05, the dataset consists of 2.8 million SNPs.

Simulation 2 has the following population split times: *t*_1_ = 200, *t*_2_ = 120, *t*_3_ = 80 and *t*_4_ = 40 with an assumed constant population size of 10,000 at all points. After filtering for minor allele frequency threshold of 0.05, the dataset consists of 2.7 million SNPs.

Simulation 3 has the following population split times: *t*_1_ = 1000, *t*_2_ = 600, *t*_3_ = 400 and *t*_4_ = 200 with an assumed constant population size of 10,000 at all points. After filtering for minor allele frequency threshold of 0.05, the dataset consists of 2.7 million SNPs.

Simulation 4 has the following population split times: *t*_1_ = 1000, *t*_2_ = 600, *t*_3_ = 400 and *t*_4_ = 200 with an assumed constant population size of 50,000 at all points. After filtering for minor allele frequency threshold of 0.05, the dataset consists of 13.4 million SNPs.

In all simulation scenarios, we have used a fixed window size of 1024 SNPs, corresponding to a mean window size of 0.37 Mb in scenario 1, 2, 3 and 0.08 Mb in scenario 4.

### 1000 Genomes Project

We have applied `HaploNet` to the phase 3 data of the 1000 Genomes Project (TGP) [1]. The entire dataset consists of phased genotype data of 2504 unrelated individuals from 26 different populations that are assigned to five super populations, which are African (AFR), Admixed American (AMR), East Asian (EAS), European (EUR) and South Asian (SAS), and we inferred local haplotype structure and global population structure for each super population. The information of the dataset and each super population is summarized in Table 2.

For each of the super populations, we have filtered their SNPs with a minor allele frequency threshold of 0.05. The AFR dataset consists of 661 individuals and 8.4 million SNPs from the following seven populations, African Caribbean in Barbados (ACB), Gambian in Western Division (GWD), Esan in Nigeria (ESN), Mende in Sierra Leone (MSL), Yoruba in Ibadan (YRI), Luhya in Webuye (LWK) and African Ancestry in Southwest US (ASW). The AMR dataset consists of 347 individuals and 6.2 million SNPs from the following four populations, Puerto Rican in Puerto Rico (PUR), Colombian in Medellin (CLM), Peruvian in Lima (PEL) and Mexican Ancestry in Los Angeles. The EAS dataset consists of 504 individuals and 5.6 million SNPs from the following five populations, Han Chinese South (CHS), Chinese Dai in Xishuangbanna (CDX), Kinh in Ho Chi Minh City (KHV), Han Chinese in Beijing (CHB) and Japanese in Tokyo (JPT). The EUR dataset consists of 503 individuals and 6 million SNPs from the following five populations, British in England and Scotland (GBR), Finnish in Finland (FIN), Iberian populations in Spain (IBS), Utah residents with Northern and Western European ancestry (CEU) and Toscani in Italy (TSI). The SAS dataset consists of 489 individuals and 6.2 million SNPs from the following seven populations, Punjabi in Lahore (PJL), Bengali in Bangladesh (BEB), Sri Lankan Tamil in the UK (STU), Indian Telugu in the UK (ITU) and Gujarati Indians in Houston (GIH). We have used a fixed window size of 1024 SNPs for all super populations.

We have additionally used the EUR super population to evaluate various aspects of our proposed method. We have applied `HaploNet` to a filtered SNP set that overlap with the SNP set of the high density genotype chip data of the 1000 Genomes project to explore our capabilities on genotype chip datasets, while using a lower window size of 256. We have applied `HaploNet` on haplotype matrices with permuted SNPs to ensure that our model captures and utilizes the LD information in the haplotypes. We have also tested different window sizes (*L* ={512, 1024, 2048}) to evaluate their effect on the inference of global population structure. Lastly, we have also tested a simpler version of our model architecture where we only use the categorical latent variable with a linear decoder to investigate the importance of the flexible Gaussian mixture prior. This version can be seen as amortized haplotype clustering where the decoder learns allele frequencies for each haplotype cluster.

The 1000 Genomes Project phase 3 data used in this study is publicly available at https://www.internationalgenome.org/category/phase-3/.

### UK Biobank

To test the scalabilty of `HaploNet`, we have also applied it to array data of the UK Biobank using unrelated individuals who are self-reported as ‘White British’ as well as having similar genetic ancestry based on PCA from genotype data. We only use SNPs from the UK Biobank Axiom array, where we have filtered the SNPs based on available QC information, a minor allele frequency threshold of 0.01, a maximum missingness rate of 0.1 and additionally removed variants in located known high LD regions. The final dataset consists of 276,732 individuals and 567,499 SNPs. We perform phasing on the genotype data using `SHAPEIT4` without using a reference panel. Further details of the sample and variant filtering are described in the supplementary material. We have used a fixed window size of 256 SNPs, corresponding to a mean window size of 0.6 Mb.

The genotype data from the UK Biobank can be obtained by application https://www.ukbiobank.ac.uk/.

### Computational comparisons

All models of `HaploNet`, as well as other analyses, presented in this study have been run on a machine with a NVIDIA GeForce RTX 2080 Ti GPU (11GB VRAM), using CUDA v.10.2 and cuDNN v.7.6.5, and a Intel Core i9-10920X CPU (3.5 GHz, 24 threads). We compare `HaploNet` with widely used software based on performance as well as runtime.

We have compared the estimated ancestry proportions of `HaploNet` with the results of the widely used software `ADMIXTURE` (v.1.3) [2], which uses unphased genotypes, in the simulation scenarios and in each of the super populations in the 1000 Genomes Project. Both methods have been run at least five times using different seeds and determined convergence with three runs being within 10 log-likelihood units of the highest achieved log-likelihood. We evaluate their performances to distinguish populations based on a signal-to-noise ratio measure using the available population labels. We use the average within-population distance of ancestry proportions in comparison to the average between-population distance as a measure of how well populations are separable, similarly to the approach in [31]. We included code and scripts for the signal-to-noise ratio measure in the GitHub repository and it is fully detailed in the supplementary material (S3).

For estimating the covariance matrix and performing PCA, we have compared `HaploNet` to `ChromoPainter` (v.4.1.0) [32] and standard PCA in `PLINK` (v.2.0) [11] on unphased genotypes. We have used `ChromoPainter` to estimate the shared genome chunks between individuals in an unsupervised manner such that no population information is given and all individuals can be used as donors of each other. We are using their linked model that utilizes pre-estimated recombination rates from genetic maps of the human chromosomes to model the correlation between SNPs using default parameters. We have used their own R library for performing PCA on the estimated chunkcounts matrix. As well as for the ancestry estimations, we also evaluate the population structure inferred using PCA based on the signal-to-noise ratio measure with the principal components. We further compare the computational runtime of `HaploNet` and `ChromoPainter` on chromosome 2 for each of the super populations.

## Acknowledgements

The study was supported by the Lundbeck foundation (R215-2015-4174). This research was conducted using the UK Biobank Resource under application 32683. JM and AA conceived the study and derived the methods. JM implemented the methods and performed the analyses. JM and AA discussed the results and contributed to the final manuscript.

## References

[1] 1000 Genomes Project Consortium et al. A global reference for human genetic variation. Nature, 526(7571):68–74, 2015.

[2] D. H. Alexander,, J. Novembre,, and K. Lange,. Fast model-based estimation of ancestry in unrelated individuals. Genome research, 19(9):1655–1664, 2009.

[3] K. Ausmees, and C. Nettelblad,. A deep learning framework for characterization of genotype data. G3, 12(3):jkac020, 2022.

[4] P. Baldi,. Autoencoders, unsupervised learning, and deep architectures. In Proceedings of ICML workshop on unsupervised and transfer learning, pages 37–49, 2012.

[5] C. Battey,, G. C. Coffing,, and A. D. Kern,. Visualizing population structure with variational autoencoders. BioRxiv, 2020.

[6] S. Behnel,, R. Bradshaw,, C. Citro,, L. Dalcin,, D. S. Seljebotn,, and K. Smith,. Cython: The best of both worlds. Computing in Science & Engineering, 13(2):31–39, 2011.

[7] D. M. Blei,, A. Kucukelbir,, and J. D. McAuliffe,. Variational inference: A review for statisticians. Journal of the American statistical Association, 112(518):859–877, 2017.

[8] D. M. Blei,, A. Kucukelbir,, and J. D. McAuliffe,. Variational inference: A review for statisticians. Journal of the American statistical Association, 112(518):859–877, 2017.

[9] Y. Bozkurt Varolgunes,, T. Bereau,, and J. F. Rudzinski,. Interpretable embeddings from molecular simulations using gaussian mixture variational autoencoders. arXiv e-prints, pages arXiv– 1912, 2019.

[10] J. Chan,, V. Perrone,, J. Spence,, P. Jenkins,, S. Mathieson,, and Y. Song,. A likelihood-free inference framework for population genetic data using exchangeable neural networks. In Advances in neural information processing systems, pages 8594–8605, 2018.

[11] C. C. Chang,, C. C. Chow,, L. C. Tellier,, S. Vattikuti,, S. M. Purcell,, and J. J. Lee,. Second-generation plink: rising to the challenge of larger and richer datasets. Gigascience, 4(1):s13742– 015, 2015.

[12] M. Collier, and H. Urdiales,. Scalable deep unsupervised clustering with concrete gmvaes. arXiv preprint 1909.08994, 2019.

[13] A. Diaz-Papkovich,, L. Anderson-Trocmé,, C. Ben-Eghan,, and S. Gravel,. Umap reveals cryptic population structure and phenotype heterogeneity in large genomic cohorts. PLoS genetics, 15(11):e1008432, 2019.

[14] N. Dilokthanakul,, P. A. Mediano,, M. Garnelo,, M. C. Lee,, H. Salimbeni,, K. Arulkumaran,, and M. Shanahan,. Deep unsupervised clustering with gaussian mixture variational autoencoders. arXiv preprint 1611.02648, 2016.

[15] C. Ding, and X. He,. K-means clustering via principal component analysis. In Proceedings of the twenty-first international conference on Machine learning, page 29, 2004.

[16] J. G. A. Figueroa,. Gaussian mixture variational autoencoder. https://github.com/jariasf/GMVAE. Accessed: 2022-01-18.

[17] L. Flagel,, Y. Brandvain,, and D. R. Schrider,. The unreasonable effectiveness of convolutional neural networks in population genetic inference. Molecular biology and evolution, 36(2):220– 238, 2019.

[18] G. R. Gower,, P. I. Picazo,, M. Fumagalli,, and F. Racimo,. Detecting adaptive introgression in human evolution using convolutional neural networks. bioRxiv, 2020.

[19] S. Gravel,, F. Zakharia,, A. Moreno-Estrada,, J. K. Byrnes,, M. Muzzio,, J. L. Rodriguez-Flores,, E. E. Kenny,, C. R. Gignoux,, B. K. Maples,, W. Guiblet,, et al. Reconstructing native american migrations from whole-genome and whole-exome data. PLoS genetics, 9(12):e1004023, 2013.

[20] B. C. Haller, and P. W. Messer,. Slim 3: Forward genetic simulations beyond the wright–fisher model. Molecular biology and evolution, 36(3):632–637, 2019.

[21] C. R. Harris,, K. J. Millman,, S. J. van der Walt,, R. Gommers,, P. Virtanen,, D. Cournapeau,, E. Wieser,, J. Taylor,, S. Berg,, N. J. Smith,, et al. Array programming with numpy. Nature, 585(7825):357–362, 2020.

[22] K. Hornik,, M. Stinchcombe,, and H. White,. Universal approximation of an unknown mapping and its derivatives using multilayer feedforward networks. Neural networks, 3(5):551–560, 1990.

[23] S. Ioffe, and C. Szegedy,. Batch normalization: Accelerating deep network training by reducing internal covariate shift. arXiv preprint 1502.03167, 2015.

[24] E. Jang,, S. Gu,, and B. Poole,. Categorical reparameterization with gumbel-softmax. arXiv preprint 1611.01144, 2016.

[25] Z. Jiang,, Y. Zheng,, H. Tan,, B. Tang,, and H. Zhou,. Variational deep embedding: An unsupervised and generative approach to clustering. arXiv preprint 1611.05148, 2016.

[26] J. Kelleher, and K. Lohse,. Coalescent simulation with msprime. Statistical Population Genomics, page 191, 2020.

[27] D. P. Kingma, and J. Ba,. Adam: A method for stochastic optimization. arXiv preprint 1412.6980, 2014.

[28] D. P. Kingma,, S. Mohamed,, D. J. Rezende,, and M. Welling,. Semi-supervised learning with deep generative models. In Advances in neural information processing systems, pages 3581– 3589, 2014.

[29] D. P. Kingma,, T. Salimans,, R. Jozefowicz,, X. Chen,, I. Sutskever,, and M. Welling,. Improved variational inference with inverse autoregressive flow. In Advances in neural information processing systems, pages 4743–4751, 2016.

[30] D. P. Kingma, and M. Welling,. Auto-encoding variational bayes. arXiv preprint 1312.6114, 2013.

[31] D. J. Lawson, and D. Falush,. Population identification using genetic data. Annual review of genomics and human genetics, 13:337–361, 2012.

[32] D. J. Lawson,, G. Hellenthal,, S. Myers,, and D. Falush,. Inference of population structure using dense haplotype data. PLoS Genet, 8(1):e1002453, 2012.

[33] N. Li, and M. Stephens,. Modeling linkage disequilibrium and identifying recombination hotspots using single-nucleotide polymorphism data. Genetics, 165(4):2213–2233, 2003.

[34] Z. Li,, A. Löytynoja,, A. Fraimout,, and J. Merilä,. comparisons. R Soc Open Sci, 6(11):190666, Nov 2019.

[35] S. Ma, and G. Shi,. On rare variants in principal component analysis of population stratification. BMC genetics, 21(1):1–11, 2020.

[36] C. J. Maddison,, A. Mnih,, and Y. W. Teh,. The concrete distribution: A continuous relaxation of discrete random variables. arXiv preprint 1611.00712, 2016.

[37] G. McVean,. A genealogical interpretation of principal components analysis. PLoS Genet, 5(10):e1000686, Oct 2009.

[38] A. Miles, and N. Harding,. scikit-allel: A python package for exploring and analysing genetic variation data, 2017.

[39] D. M. Montserrat,, C. Bustamante,, and A. Ioannidis,. Class-conditional vae-gan for local-ancestry simulation. arXiv preprint 1911.13220, 2019.

[40] A. Paszke,, S. Gross,, F. Massa,, A. Lerer,, J. Bradbury,, G. Chanan,, T. Killeen,, Z. Lin,, N. Gimelshein,, L. Antiga,, et al. Pytorch: An imperative style, high-performance deep learning library. In Advances in neural information processing systems, pages 8026–8037, 2019.

[41] N. Patterson,, A. L. Price,, and D. Reich,. Population structure and eigenanalysis. PLoS genet, 2(12):e190, 2006.

[42] J. K. Pritchard,, M. Stephens,, and P. Donnelly,. Inference of population structure using multilocus genotype data. Genetics, 155(2):945–959, 2000.

[43] F. Privé,, K. Luu,, M. G. Blum,, J. J. McGrath,, and B.J. Vilhjálmsson,. Efficient toolkit implementing best practices for principal component analysis of population genetic data. Bioinformatics, 36(16):4449–4457, 2020.

[44] D. J. Rezende, and S. Mohamed,. Variational inference with normalizing flows. arXiv preprint 1505.05770, 2015.

[45] D. J. Rezende,, S. Mohamed,, and D. Wierstra,. Stochastic backpropagation and approximate inference in deep generative models. arXiv preprint 1401.4082, 2014.

[46] D. E. Rumelhart,, G. E. Hinton,, and R. J. Williams,. Learning internal representations by error propagation. Technical report, California Univ San Diego La Jolla Inst for Cognitive Science, 1985.

[47] J. N. Saada,, G. Kalantzis,, D. Shyr,, F. Cooper,, M. Robinson,, A. Gusev,, and P. F. Palamara,. Identity-by-descent detection across 487,409 british samples reveals fine scale population structure and ultra-rare variant associations. Nature Communications, 11, 2020.

[48] A. Saxena,, M. Prasad,, A. Gupta,, N. Bharill,, O. P. Patel,, A. Tiwari,, M. J. Er,, W. Ding,, and C.-T. Lin,. A review of clustering techniques and developments. Neurocomputing, 267:664–681, 2017.

[49] D. R. Schrider, and A. D. Kern,. Supervised machine learning for population genetics: a new paradigm. Trends in Genetics, 34(4):301–312, 2018.

[50] S. Sheehan, and Y. S. Song,. Deep learning for population genetic inference. PLoS computational biology, 12(3):e1004845, 2016.

[51] R. Shu,. Gaussian mixture vae: Lessons in variational inference, generative models, and deep nets. http://ruishu.io/2016/12/25/gmvae/. Accessed: 2022-01-18.

[52] R. Shu,, H. H. Bui,, S. Zhao,, M. J. Kochenderfer,, and S. Ermon,. Amortized inference regularization. Advances in Neural Information Processing Systems, 31:4393–4402, 2018.

[53] L. Skotte,, T. S. Korneliussen,, and A. Albrechtsen,. Estimating individual admixture proportions from next generation sequencing data. Genetics, 195(3):693–702, 2013.

[54] C. K. Sønderby,, T. Raiko,, L. Maaløe,, S. K. Sønderby,, and O. Winther,. Ladder variational autoencoders. In Advances in neural information processing systems, pages 3738–3746, 2016.

[55] H. Tang,, J. Peng,, P. Wang,, and N. J. Risch,. Estimation of individual admixture: analytical and study design considerations. Genetic Epidemiology: The Official Publication of the International Genetic Epidemiology Society, 28(4):289–301, 2005.

[56] J. A. Tennessen,, A. W. Bigham,, T. D. O’Connor,, W. Fu,, E. E. Kenny,, S. Gravel,, S. McGee,, R. Do,, X. Liu,, G. Jun,, et al. Evolution and functional impact of rare coding variation from deep sequencing of human exomes. science, 337(6090):64–69, 2012.

[57] R. Varadhan, and C. Roland,. Simple and globally convergent methods for accelerating the convergence of any em algorithm. Scandinavian Journal of Statistics, 35(2):335–353, 2008.

[58] Z. Wang,, J. Wang,, M. Kourakos,, N. Hoang,, H. H. Lee,, I. Mathieson,, and S. Mathieson,. Automatic inference of demographic parameters using generative adversarial networks. bioRxiv, 2020.

[59] J. Xie,, R. Girshick,, and A. Farhadi,. Unsupervised deep embedding for clustering analysis. In International conference on machine learning, pages 478–487, 2016.

[60] B. Yang,, X. Fu,, N. D. Sidiropoulos,, and M. Hong,. Towards k-means-friendly spaces: Simultaneous deep learning and clustering. In international conference on machine learning, pages 3861–3870. PMLR, 2017.

[61] B. Yelmen,, A. Decelle,, L. Ongaro,, D. Marnetto,, F. Montinaro,, C. Furtlehner,, L. Pagani,, and F. Jay,. Creating artificial human genomes using generative models. 2019.

[62] A. Zimek,, E. Schubert,, and H.-P. Kriegel,. A survey on unsupervised outlier detection in highdimensional numerical data. Statistical Analysis and Data Mining: The ASA Data Science Journal, 5(5):363–387, 2012.

