## Supplementary Material for "Haplotype and Population Structure Inference using Neural Networks in Whole-Genome Sequencing Data"

#### S1 Derivation of the evidence lower bound (ELBO)

We here derive the ELBO from the marginal log-likelihood of a haplotype, $\mathbf{x}$, using Jensen's inequality and our following definitions, $p_\theta(\mathbf{x}, \mathbf{z}, y) = p_\theta(\mathbf{x} \mid \mathbf{z}) p_\theta(\mathbf{z} \mid y) p(y)$, $q_\phi(\mathbf{z}, y \mid \mathbf{x}) = q_\phi(\mathbf{z} \mid \mathbf{x}, y) q_\phi(y \mid \mathbf{x})$. We follow the mathematical notation used in the machine learning literature and Kingma and Welling [6].

$$\log p_\theta(\mathbf{x}) = \log \sum_y \int_{\mathbf{z}} p_\theta(\mathbf{x}, \mathbf{z}, y)\, d\mathbf{z} \tag{S1}$$

$$= \log \sum_y \int_{\mathbf{z}} p_\theta(\mathbf{x}, \mathbf{z}, y) \frac{q_\phi(\mathbf{z}, y \mid \mathbf{x})}{q_\phi(\mathbf{z}, y \mid \mathbf{x})}\, d\mathbf{z} \tag{S2}$$

$$= \log \mathbb{E}_{q_\phi(\mathbf{z}, y \mid \mathbf{x})} \left[ \frac{p_\theta(\mathbf{x}, \mathbf{z}, y)}{q_\phi(\mathbf{z}, y \mid \mathbf{x})} \right] \tag{S3}$$

$$\geq \mathbb{E}_{q_\phi(\mathbf{z}, y \mid \mathbf{x})} \left[ \log \frac{p_\theta(\mathbf{x}, \mathbf{z}, y)}{q_\phi(\mathbf{z}, y \mid \mathbf{x})} \right] \tag{S4}$$

$$= \mathbb{E}_{q_\phi(\mathbf{z}, y \mid \mathbf{x})} \left[ \log \frac{p_\theta(\mathbf{x} \mid \mathbf{z}) p_\theta(\mathbf{z} \mid y) p(y)}{q_\phi(\mathbf{z} \mid \mathbf{x}, y) q_\phi(y \mid \mathbf{x})} \right] \tag{S5}$$

$$= \mathbb{E}_{q_\phi(\mathbf{z}, y \mid \mathbf{x})} \left[ \log p_\theta(\mathbf{x} \mid \mathbf{z}) - \log \frac{q_\phi(\mathbf{z} \mid \mathbf{x}, y)}{p_\theta(\mathbf{z} \mid y)} - \log \frac{q_\phi(y \mid \mathbf{x})}{p(y)} \right]. \tag{S6}$$

We can rewrite the last term in the expectation using $p(y = c) = C^{-1}$, for $c = 1, \ldots, C$:

$$\mathbb{E}_{q_\phi(\mathbf{z}, y \mid \mathbf{x})} \left[ \log \frac{q_\phi(y \mid \mathbf{x})}{p(y)} \right] = \mathbb{E}_{q_\phi(y \mid \mathbf{x})} \mathbb{E}_{q_\phi(\mathbf{z} \mid \mathbf{x}, y)} \left[ \log \frac{q_\phi(y \mid \mathbf{x})}{p(y)} \right] \tag{S7}$$

$$= \mathbb{E}_{q_\phi(y \mid \mathbf{x})} \left[ \log \frac{q_\phi(y \mid \mathbf{x})}{p(y)} \right] \tag{S8}$$

$$= \mathbb{E}_{q_\phi(y \mid \mathbf{x})} \left[ \log q_\phi(y \mid \mathbf{x}) \right] - \mathbb{E}_{q_\phi(y \mid \mathbf{x})} \left[ \log p(y) \right] \tag{S9}$$

$$= -H(q_\phi(y \mid \mathbf{x})) + \log C, \tag{S10}$$

where $H(X)$ is the entropy of a given distribution. In that way, both the second and third term will act as regularization on the reconstruction term through the latent space distributed by $q_\phi(\mathbf{z}, y \mid \mathbf{x})$.

#### S2 Derivation of EM algorithm

Here we derive the EM algorithm for estimating ancestry proportions $\mathbf{Q} \in [0,1]^{N\times K}$ and window-based haplotype frequencies $\mathbf{F} \in [0,1]^{W\times K\times C}$, and we will use the following notation:

$W$ number of windows

$N$ number of individuals

$C$ number of haplotype states

$K$ number of ancestral states

$\mathbf{x}_{i,a}^{(w)}$ observed data of the $a$-th haplotype of individual $i$ in window $w$

$\psi^{(n)}$ auxiliary variable in iteration $n$

$q_{ik}^{(n)}$ ancestry proportion of state $k$ for individual $i$ in iteration $n$

$f_{wkc}^{(n)}$ ancestral frequency of haplotype state $c$ in ancestral state $k$ in window $w$ in iteration $n$

We define the likelihood function:

$$\mathcal{L}(\mathbf{Q},\mathbf{F};\mathbf{X}) \propto \prod_{w=1}^{W}\prod_{i=1}^{N}\prod_{a=1}^{2} p(\mathbf{x}_{i,a}^{(w)} \mid \psi) \tag{S11}$$

$$p(\mathbf{x}_{i,a}^{(w)} \mid \psi) = \sum_{k=1}^{K}\sum_{c=1}^{C} p(\mathbf{x}_{i,a}^{(w)} \mid \psi, k, y=c)p(y=c \mid k,\psi)p(k \mid \psi) \tag{S12}$$

$$= \sum_{k=1}^{K}\sum_{c=1}^{C} p(\mathbf{x}_{i,a}^{(w)} \mid y=c) f_{wkc} q_{ik}. \tag{S13}$$

The EM update for $q_{ik}$ is defined as follows for iteration $n+1$:

$$q_{ik}^{(n+1)} = \frac{\sum_{w=1}^{W}\sum_{a=1}^{2} p(k \mid \mathbf{x}_{i,a}^{(w)}, \psi^{(n)})}{\sum_{k'=1}^{K}\sum_{w=1}^{W}\sum_{a=1}^{2} p(k' \mid \mathbf{x}_{i,a}^{(w)}, \psi^{(n)})} \tag{S14}$$

$$= \frac{\sum_{w=1}^{W}\sum_{a=1}^{2} p(k \mid \mathbf{x}_{i,a}^{(w)}, \psi^{(n)})}{2W}, \tag{S15}$$

and the update for $f_{wkc}$:

$$f_{wkc}^{(n+1)} = \frac{\sum_{i=1}^{N}\sum_{a=1}^{2} p(k, y=c \mid \mathbf{x}_{i,a}^{(w)}, \psi^{(n)})}{\sum_{c'=1}^{C}\sum_{i=1}^{N}\sum_{a=1}^{2} p(k, y=c' \mid \mathbf{x}_{i,a}^{(w)}, \psi^{(n)})}, \tag{S16}$$

where

$$p(k \,|\, \mathbf{x}_{i,a}^{(w)}, \psi^{(n)}) = \sum_{c=1}^{C} p(k, y = c \,|\, \mathbf{x}_{i,a}^{(w)}, \psi^{(n)}), \tag{S17}$$

and

$$\begin{aligned}
p(k, y = c \,|\, \mathbf{x}_{i,a}^{(w)}, \psi^{(n)}) &= \frac{p(\mathbf{x}_{i,a}^{(w)} \,|\, k, y = c, \psi^{(n)}) p(y = c, \mathbf{x}_{i,a}^{(w)} \,|\, \psi^{(n)})}{\sum_{k'=1}^{K} \sum_{c'=1}^{C} p(\mathbf{x}_{i,a}^{(w)} \,|\, k', y = c', \psi^{(n)}) p(y = c', \mathbf{x}_{i,a}^{(w)} \,|\, \psi^{(n)})} && \text{(S18)} \\
&= \frac{p(\mathbf{x}_{i,a}^{(w)} \,|\, y = c) p(y = c \,|\, k, \psi^{(n)}) p(k \,|\, \psi^{(n)})}{\sum_{k'=1}^{K} \sum_{c'=1}^{C} p(\mathbf{x}_{i,a}^{(w)} \,|\, y = c') p(y = c' \,|\, k', \psi^{(n)}) p(k' \,|\, \psi^{(n)})} && \text{(S19)} \\
&= \frac{p(\mathbf{x}_{i,a}^{(w)} \,|\, y = c) f_{wkc}^{(n)} q_{ik}^{(n)}}{\sum_{k'=1}^{K} \sum_{c'=1}^{C} p(\mathbf{x}_{i,a}^{(w)} \,|\, y = c') f_{wk'c'}^{(n)} q_{ik'}^{(n)}}. && \text{(S20)}
\end{aligned}$$

### S3 Signal-to-noise ratio measure

Here we define the signal-to-noise ratio measure used to quantify how separable populations are given their labels. It is defined as the mean between-population distance relative to the mean within-population distance of a given measure,

$$\frac{\sum_p^P \frac{1}{N_{p,b}} \sum_i^N \sum_j^N \mathbb{I}[p_i \neq p_j] d_{ij}}{\sum_p^P \frac{1}{N_{p,w}} \sum_i^N \sum_j^N \mathbb{I}[p_i = p_j] d_{ij}} \tag{S21}$$

where $p$ is a population label, $P$ is the total number of population labels, $p_i$ is the population label of the $i$-th individual and $d_{ij}$ is the euclidean distance of individual $i$ and $j$ given either their ancestry proportions or principal components. $N_{p,b}$ is the number of comparisons where the population labels does not match (between) and $N_{p,w}$ is the number of comparisons where population labels match for population $p$ (within).

#### S4 Reparameterization tricks

End-to-end learning is a desired trait in deep learning models as it enables joint training of model parameters through backpropagation. However with the network architectures of variational autoencoders, the sampling steps of the latent variables are not letting information flow end-to-end as they are non-differentiable. The reparameterization trick is a simple approach to express a random variable as deterministic using the introduction of an auxiliary variable as defined in the SGVB estimator [6]. In this way, the stochastic process of sampling is removed from the computation graph and we can therefore efficiently estimate gradients and perform end-to-end learning of our network parameters, $(\phi, \theta)$, through Monte Carlo samples of the latent variables to approximate the ELBO in Equation S1.

A $D$-dimensional sample from a multivariate Gaussian distribution, $\mathbf{z}$, can be expressed using the introduction of auxiliary noise variable $\boldsymbol{\epsilon}$, with $\epsilon_d \sim \mathcal{N}(0, 1)$, for $d = 1, \ldots, D$:

$$z_d = \mu_d + \sigma_d \epsilon_d. \tag{S22}$$

Similarly, the reparameterization trick for the Gumbel-Softmax distribution is an approach for providing differentiable samples that approximate samples from a categorical distribution. It is an extension of the Gumbel-Max trick [8] for sampling from a categorical distribution, where the softmax function is used as a continuous differentiable approximation to the arg max function. The Gumbel-Softmax distribution was derived simultaneously in two independent studies [4, 7]. A $C$-dimensional one-hot encoded categorical sample $y$ can be approximated as sample $\tilde{y}$ from the Gumbel-Softmax distribution, and it is defined as follows for its entry $c$:

$$\tilde{y}_c = \frac{\exp((\log(\pi_c) + g_c)/\tau)}{\sum_{c'=1}^{C} \exp((\log(\pi_{c'}) + g_{c'})/\tau)}. \tag{S23}$$

Here $g_c, \ldots, g_C$ are i.i.d. samples from Gumbel$(0, 1)$ (auxiliary variables) and $\pi_c$ is the probability of haplotype cluster $c$. The temperature parameter, $\tau$, is governing the smoothness of the distribution such that when $\tau$ approaches 0, samples become identical to samples from the categorical distribution. We have provided examples of the Gumbel-Softmax distribution in the next section to provide the reader with a better intuition of it. Note that $\tilde{y} \in [0, 1]^C$ unlike $y$, and the functions $\mu_\theta, \sigma_\theta, \mu_\phi, \sigma_\phi$ parameterized by the neural networks are therefore redefined as mapping to or from a continuous space in the practical implementation.

### S5 Gumbel-Softmax sampling

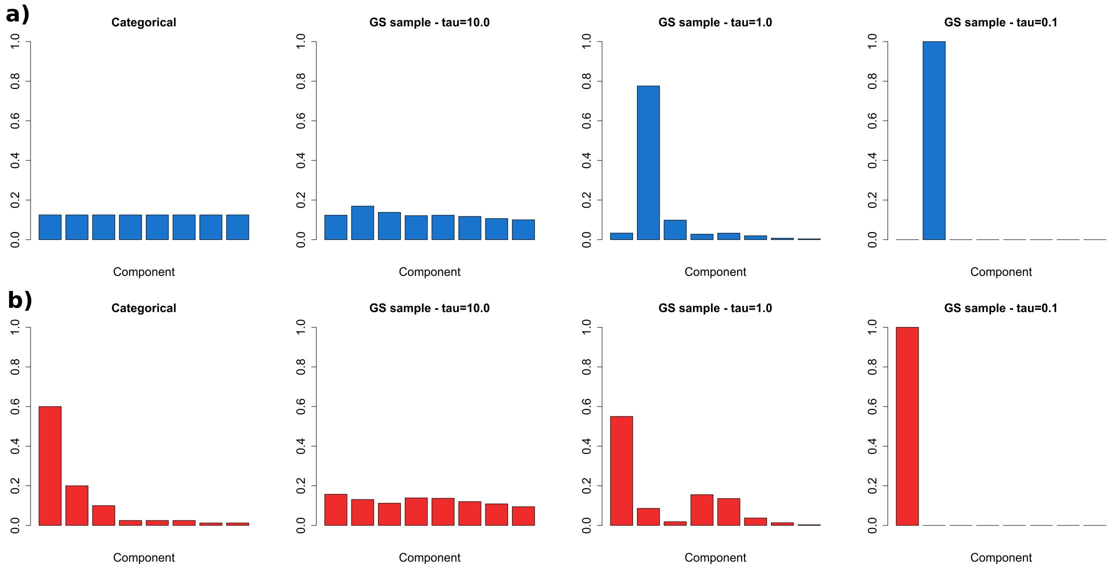


**Figure S1:** Two examples of approximate sampling from different categorical distributions using the Gumbel-Softmax distribution. First column displays the two categorical distributions with the two examples distinguishable by color, and the next three columns display random samplings from the distributions for different temperatures. The random sampling is performed using the reparameterization trick as described in Equation S23 with $\tau = 10, 1, 0.1$. ***a)*** A discrete uniform distribution, ***b)*** a categorical distribution with decreasing probabilities. Here it can be seen that as $\tau$ approaches 0, the sample from the Gumbel-Softmax distribution will converge to a categorical sample, while as $\tau$ approaches $\infty$, it will converge towards a uniform sample.

#### S6 Network architectures

In all models, the architectures for the networks of `HaploNet` were defined as follows:

- $\pi_\phi(\mathbf{x})$: $L$-$H$-ReLU-BN-$C$
- $\mu_\phi(\mathbf{x}, y)$, $\sigma^2_\phi(\mathbf{x}, y)$: $(L + C)$-$H$-ReLU-BN-$D$
- $\mu_\theta(y)$, $\sigma^2_\theta(y)$: $C$-$D$
- $\pi_\theta(\mathbf{z})$: $D$-$H$-ReLU-BN-$L$

with a hidden layer of 256 units ($H = 256$), ReLU being a ReLU activation layer, and BN a batch normalization layer.

##### Simulations

- Dimensions: $L = 1024$, $D = 64$, $C = 32$
- Batch size: 128
- Epochs: 200

##### 1000 Genomes Project (AFR, AMR, EAS, EUR, SAS) [1]

- Dimensions: $L = 1024$ [1], $D = 64$, $C = 32$
- Batch size: 128
- Epochs: 200

##### UK Biobank [3]

- Dimensions: $L = 256$, $D = 64$, $C = 32$
- Batch size: 8192
- Epochs: 100

[1] $L = 512$ and $L = 2048$ were used for evaluating the effect of window size for the EUR super population, while $L = 256$ was used for the EUR super population when only using sites on the high density genotype chip.

### S7 Supplementary figures and tables

#### S7.1 Simulations in `msprime`

Four different simulation scenarios have been modelled using `msprime` [5] for a sequence length of $1.0 \times 10^9$. Chromosomes of 100 diploid samples have been sampled from each of the five populations. A uniform recombination rate of $1.0 \times 10^{-9}$ has been assumed in all simulations as well as a mutation rate of $2.36 \times 10^{-8}$. The following population split times (in generations) have been used:

- Simulation 1: $\{t_1 = 100, t_2 = 60, t_3 = 40, t_4 = 20\}$, all population sizes: 10000.
- Simulation 2: $\{t_1 = 200, t_2 = 120, t_3 = 80, t_4 = 40\}$, all population sizes: 10000.
- Simulation 3: $\{t_1 = 1000, t_2 = 600, t_3 = 400, t_4 = 200\}$, all population sizes: 10000.
- Simulation 4: $\{t_1 = 1000, t_2 = 600, t_3 = 400, t_4 = 200\}$, all population sizes: 50000.

##### S7.1.1 Comparisons

| | `HaploNet` | `ADMIXTURE` (LD-pruned) |
|---|---|---|
| **Simulation 1** | 22.54466 | 7.86155 (8.40691) |
| **Simulation 2** | 5.84812 | 3.79932 (4.37785) |
| **Simulation 3** | 1106.03398 | 63.96098 (86.78266) |
| **Simulation 4** | 311.18541 | 16.28952 (17.86109) |

**Table S1:** Mean between-population distance relative to the mean within-population distance of estimated ancestry proportions for the simulated scenarios. A higher value means a higher signal-to-noise ratio, thus more separable populations. The measures of `ADMIXTURE` from the LD pruned datasets are also reported for the used $K$ in each super population. (LD pruning performed in `PLINK: --indep-pairwise 50 10 0.5`)

| | `HaploNet` | `ChromoPainter` | `PLINK` (LD-pruned) |
|---|---|---|---|
| **Simulation 1** | 3.79448 | 12.68871 | 3.48961 (3.69830) |
| **Simulation 2** | 7.53089 | 20.36589 | 7.60278 (8.15770) |
| **Simulation 3** | 26.66899 | 26.14253 | 32.51606 (37.17452) |
| **Simulation 4** | 16.92876 | N/A | 17.06983 (18.57127) |

**Table S2:** Mean between-population distance relative to the mean within-population distance of the inferred principal components for the simulated scenarios. Four principal components have been used in all cases. A higher value means a higher signal-to-noise ratio, thus more separable populations. The measures of the inferred principal components from the LD pruned datasets using `PLINK` is reported in parentheses (LD pruning performed in `PLINK: --indep-pairwise 50 10 0.5`). `ChromoPainter` was terminated in simulation 4 due to excessive memory requirements and estimated computational runtime.

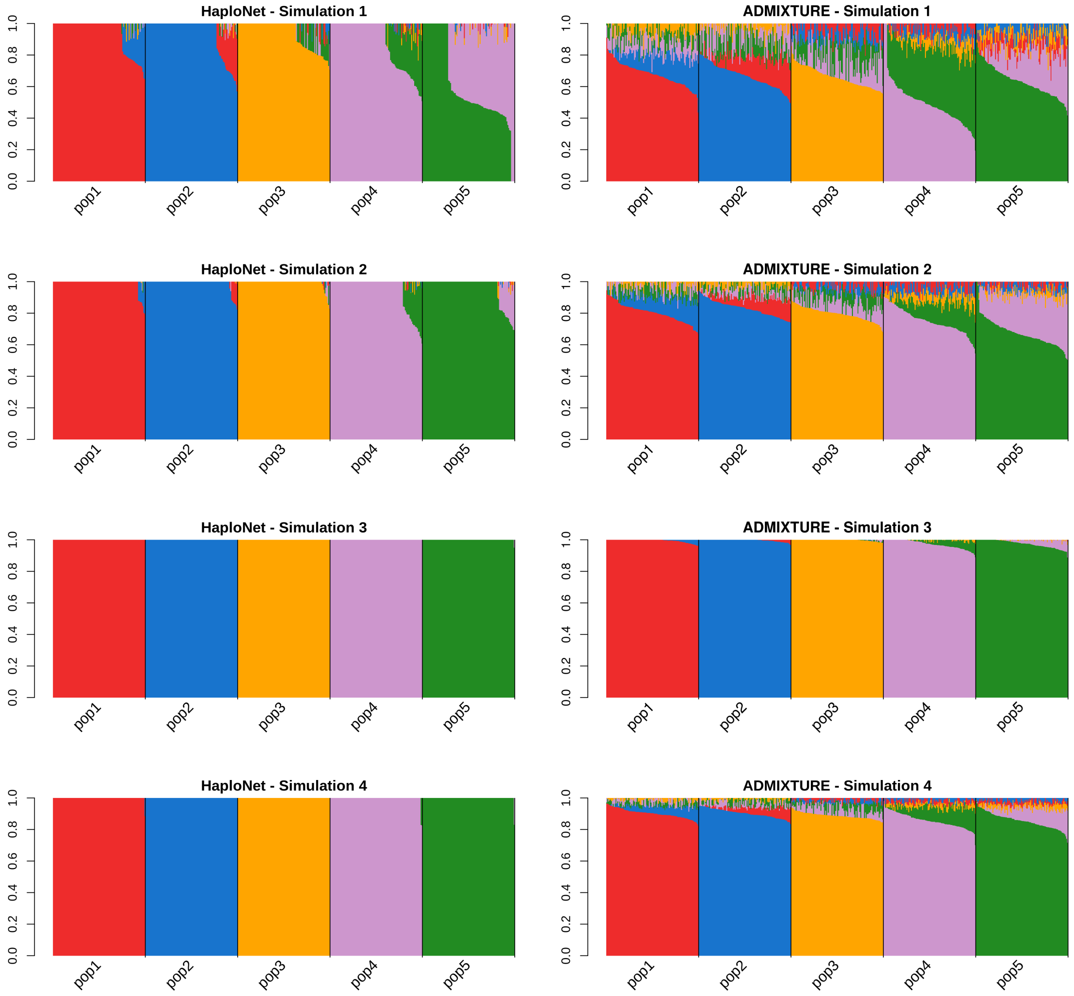


**Figure S2:** Estimated ancestry proportions in the different simulation scenarios using `HaploNet` (left column) and `ADMIXTURE` [2] (right column).

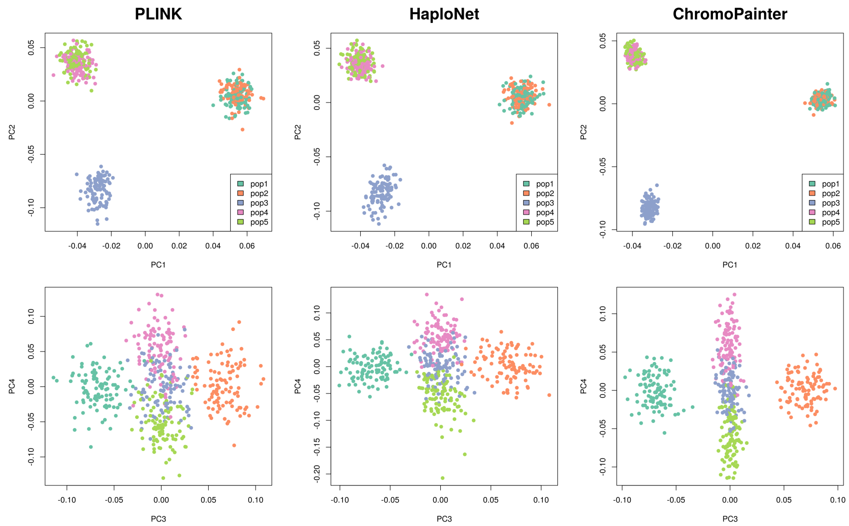


**Figure S3:** Inferred population structure in Simulation 1 based on PCA using `PLINK` (left column), `HaploNet` (middle column), and `ChromoPainter` (right column). First row displays PC1 plotted against PC2, while the second row shows PC3 against PC4.

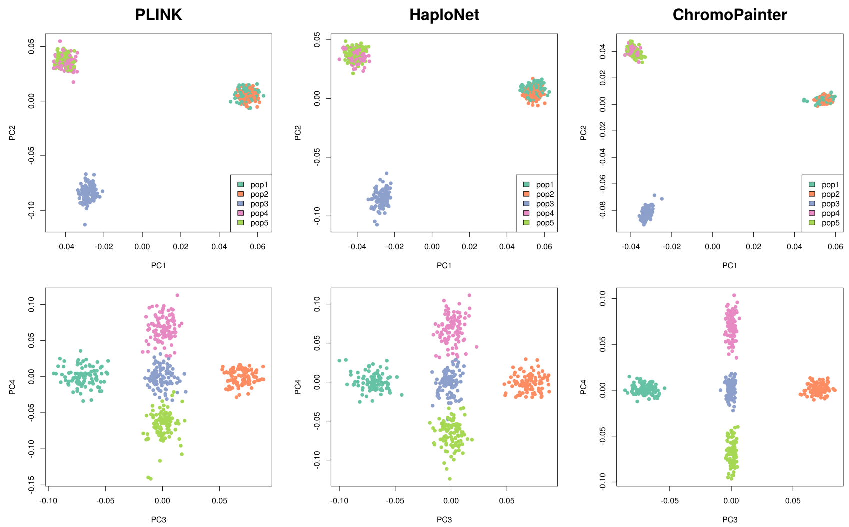


**Figure S4:** Inferred population structure in Simulation 2 based on PCA using `PLINK` (left column), `HaploNet` (middle column), and `ChromoPainter` (right column). First row displays PC1 plotted against PC2, while the second row shows PC3 against PC4.

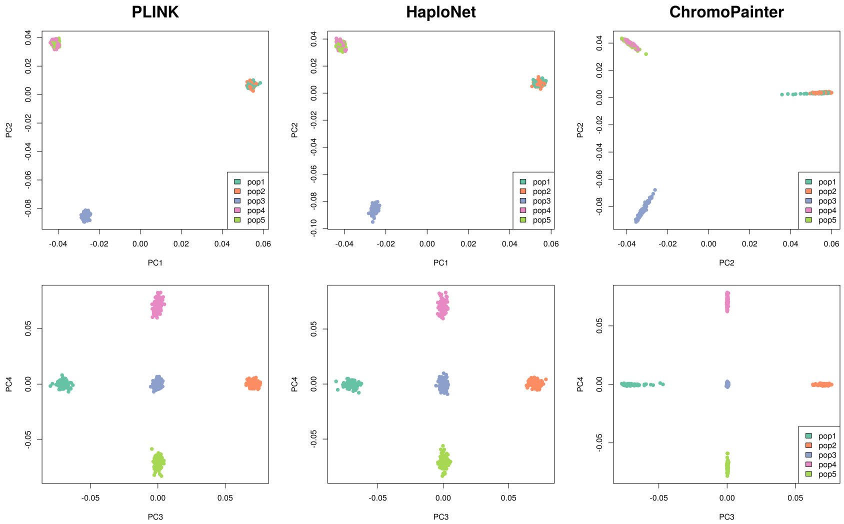


**Figure S5:** Inferred population structure in Simulation 3 based on PCA using `PLINK` (left column), `HaploNet` (middle column), and `ChromoPainter` (right column). First row displays PC1 plotted against PC2, while the second row shows PC3 against PC4.

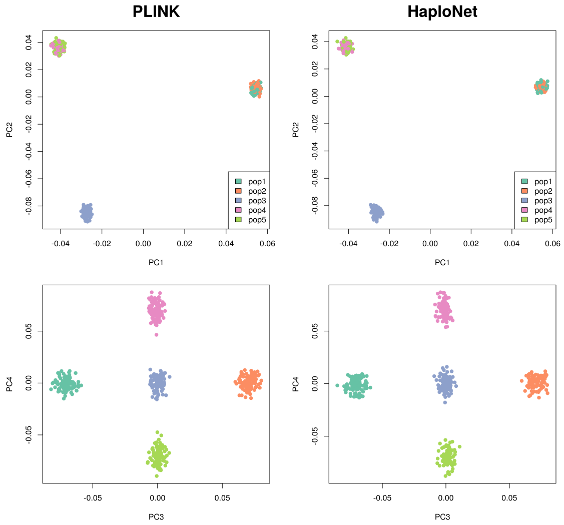


**Figure S6:** Inferred population structure in Simulation 4 based on PCA using `PLINK` (left column) and `HaploNet` (right column)). First row displays PC1 plotted against PC2, while the second row shows PC3 against PC4. `ChromoPainter` was terminated in simulation 4 due to excessive memory requirements and estimated computational runtime.

#### S7.2 1000 Genomes Project

##### S7.2.1 Comparisons

| | K=2 | K=3 | K=4 | K=5 |
|---|---|---|---|---|
| | | **AFR** | | |
| HaploNet | 5.72604 | 7.38230 | 7.30438 | 7.01651 |
| ADMIXTURE (LD-pruned) | 4.05180 | 5.45061 | 6.13836 | 4.87877 (5.84965) |
| | | **AMR** | | |
| HaploNet | 3.02893 | 2.64753 | 2.61632 | - |
| ADMIXTURE (LD-pruned) | 2.80201 | 2.13841 | 2.28056 (2.58715) | - |
| | | **EAS** | | |
| HaploNet | 8.98795 | 8.21613 | 8.05040 | - |
| ADMIXTURE (LD-pruned) | 7.41574 | 5.98761 | 3.21532 (3.31927) | - |
| | | **EUR** | | |
| HaploNet | 12.64427 | 9.80112 | 7.86159 | - |
| ADMIXTURE(LD-pruned) | 7.74596 | 5.04840 | 1.86814 (4.36921) | - |
| | | **SAS** | | |
| HaploNet | 2.82519 | 2.87521 | - | - |
| ADMIXTURE (LD-pruned) | 2.23994 | 2.06078 (2.32379) | - | - |

**Table S3:** Mean between-population distance relative to the mean within-population distance of estimated ancestry proportions for each super population of the 1000 Genomes Project. A higher value means a higher signal-to-noise ratio, thus more separable populations, while the measures between the different values of K are not comparable. The scores of `ADMIXTURE` on the LD pruned datasets are also reported for the highest used $K$ in each super population (LD pruning performed in `PLINK: --indep-pairwise 50 10 0.5`).

| | HaploNet | ChromoPainter | PLINK (LD-pruned) |
|---|---|---|---|
| **AFR** (3) | 4.40707 | 5.24017 | 4.25830 (4.55877) |
| **AMR** (4) | 2.42171 | 2.37184 | 2.37600 (2.40783) |
| **EAS** (3) | 4.36325 | 6.94765 | 3.02060 (3.42440) |
| **EUR** (3) | 3.93642 | 6.51211 | 2.55128 (4.28249) |
| **SAS** (3) | 1.78487 | 2.23910 | 1.78416 (1.92694) |

**Table S4:** Mean between-population distance relative to the mean within-population distance of the inferred principal components for each of the super populations of the 1000 Genomes Project. The number of principal components used in each scenario is reported in the parentheses. The measures of inferred principal components from the LD pruned datasets using `PLINK` is reported in parentheses (LD pruning performed in `PLINK: --indep-pairwise 50 10 0.5`).

##### S7.2.2 AFR

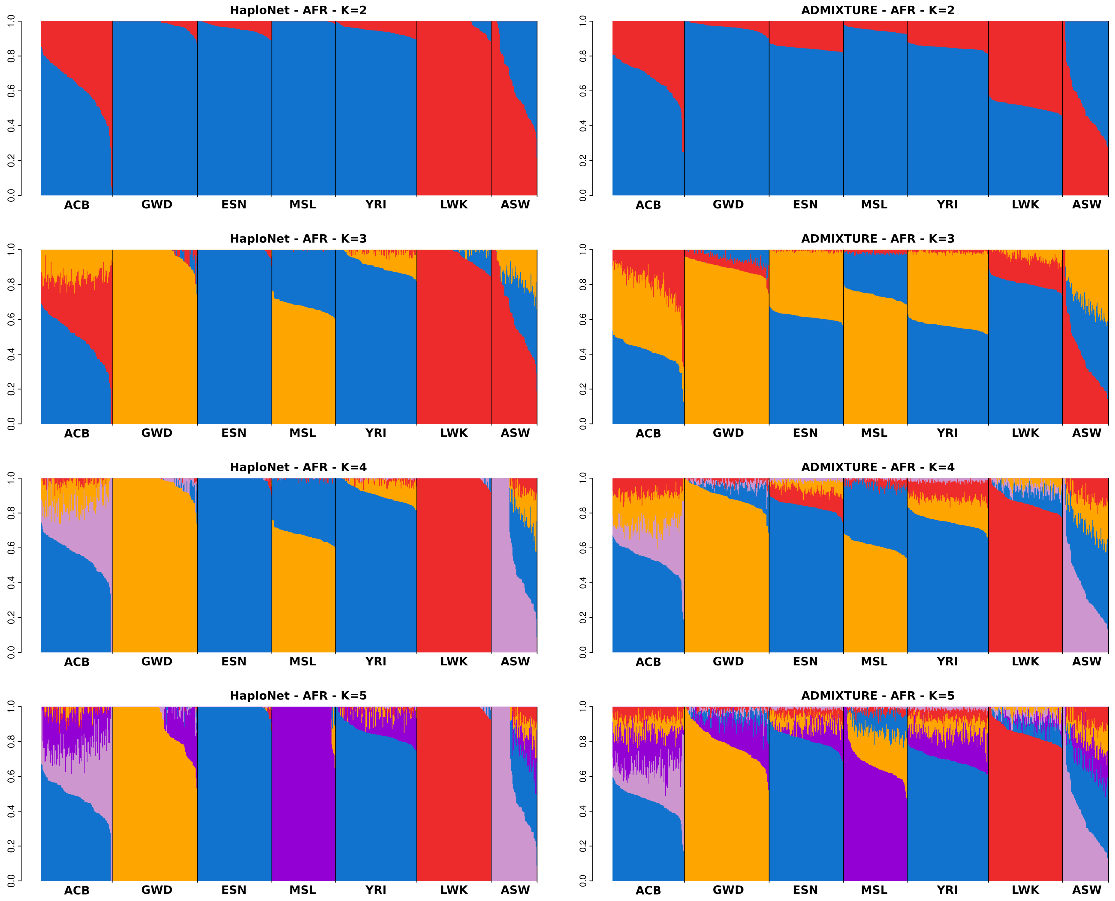


**Figure S7:** Estimated ancestry proportions in the AFR super population of the 1000 Genomes Project using `HaploNet` (left column) and `ADMIXTURE` (right column) for $K = 2, 3, 4, 5$.

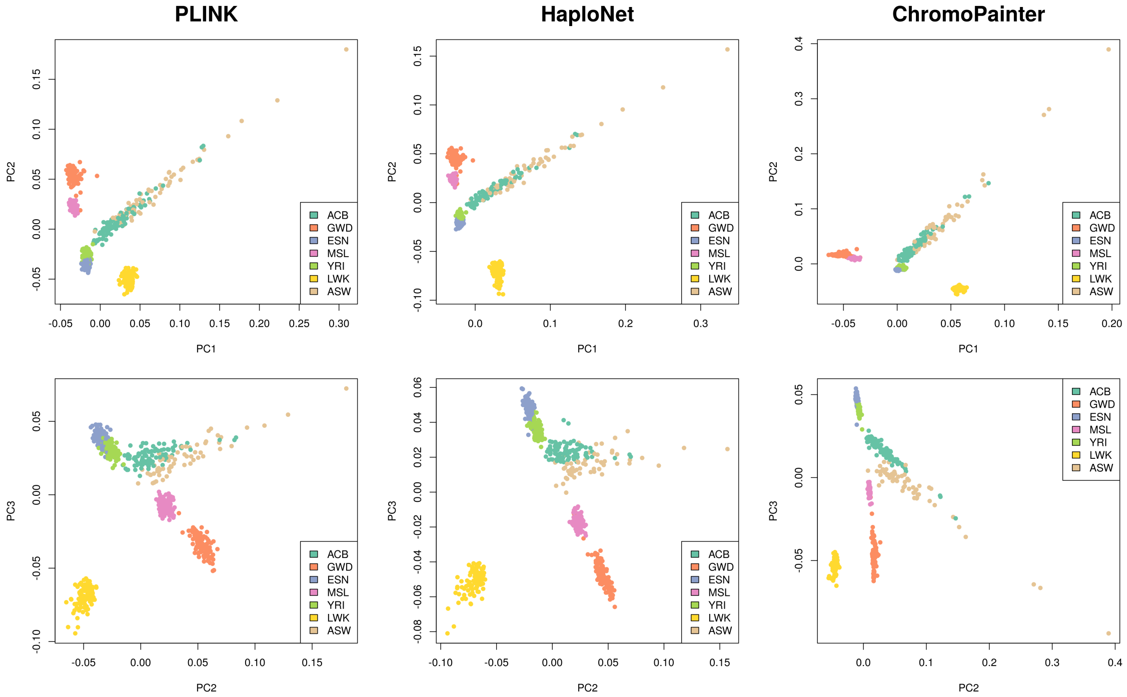


**Figure S8:** Inferred population structure in the AFR super population based on PCA using `HaploNet` (left column), `PLINK` (middle column) and `ChromoPainter` (right column). First row displays PC1 plotted against PC2, while the second row shows PC2 against PC3.

##### S7.2.3 AMR

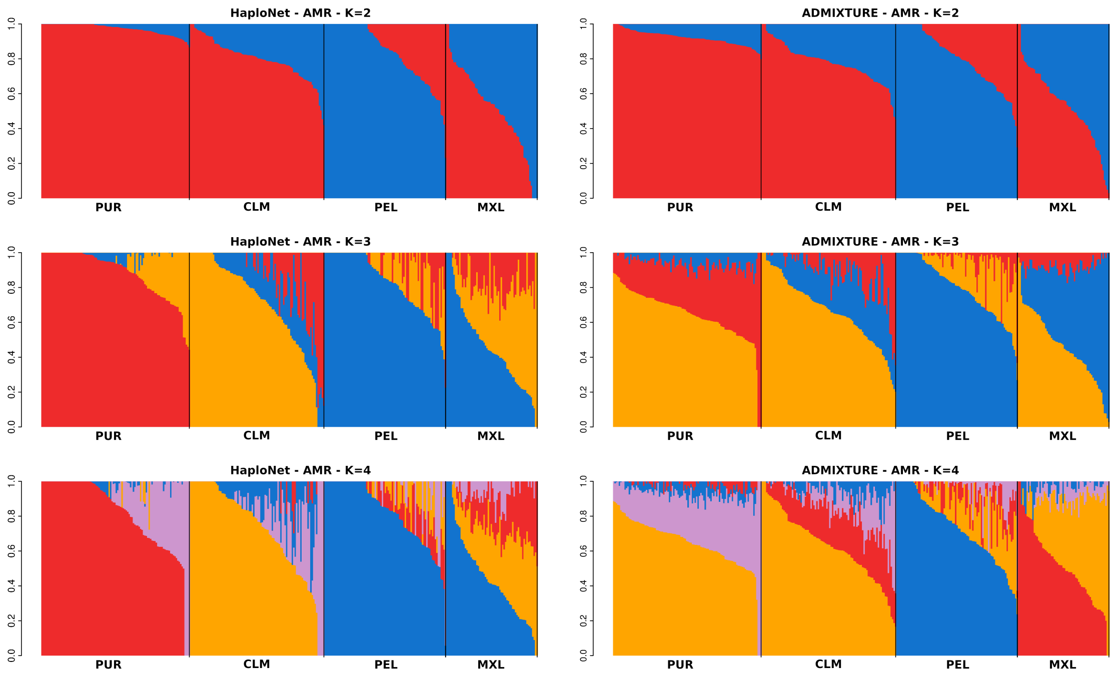


**Figure S9:** Estimated ancestry proportions in the AMR super population of the 1000 Genomes Project using `HaploNet` (left column) and `ADMIXTURE` (right column) for $K = 2, 3, 4$.

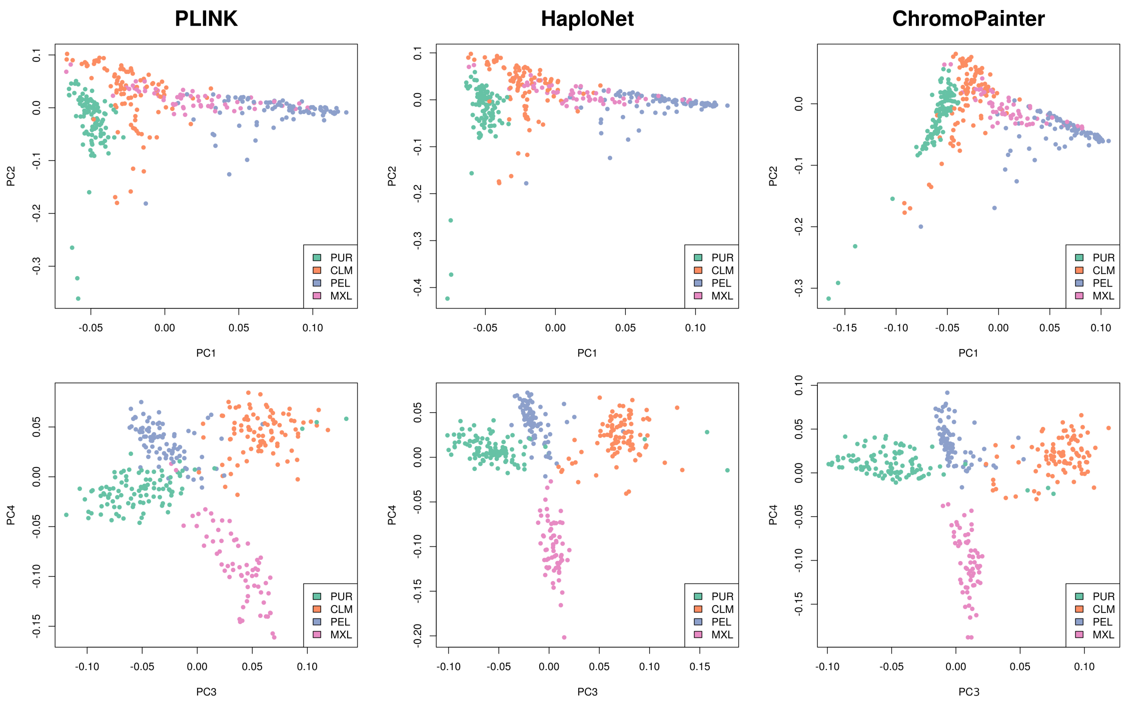


**Figure S10:** Inferred population structure in the AMR super population based on PCA using `PLINK` (left column), `HaploNet` (middle column), and `ChromoPainter` (right column). First row displays PC1 plotted against PC2, while the second row shows PC3 against PC4.

##### S7.2.4 EAS

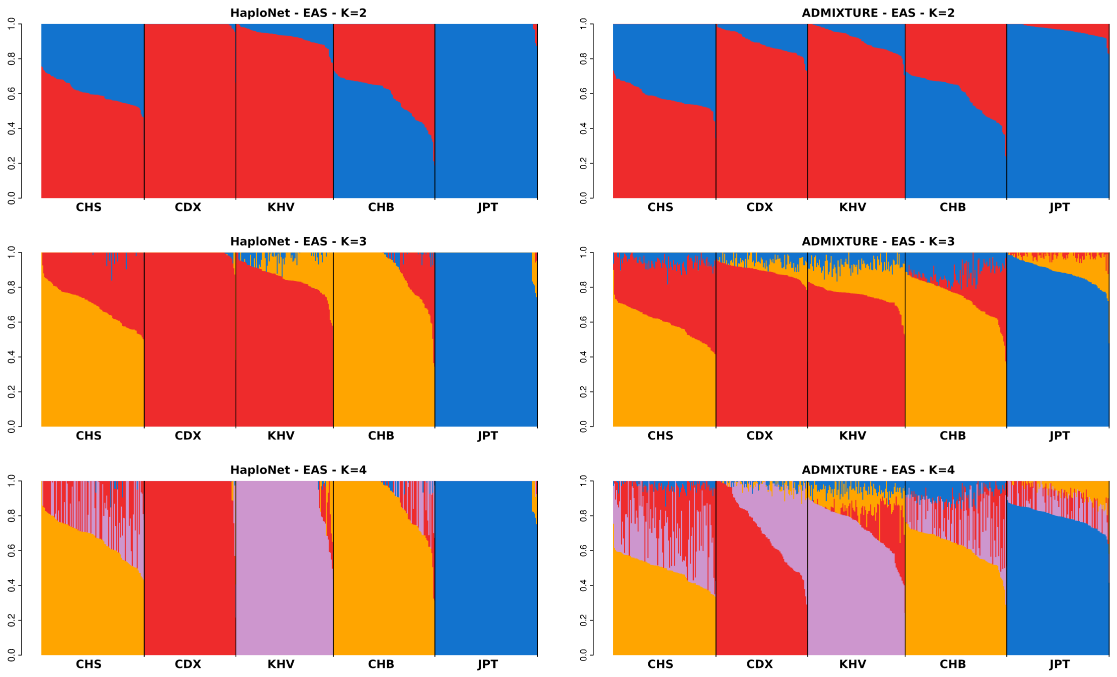


**Figure S11:** Estimated ancestry proportions in the EAS super population of the 1000 Genomes Project using `HaploNet` (left column) and `ADMIXTURE` (right column) for $K = 2, 3, 4$.

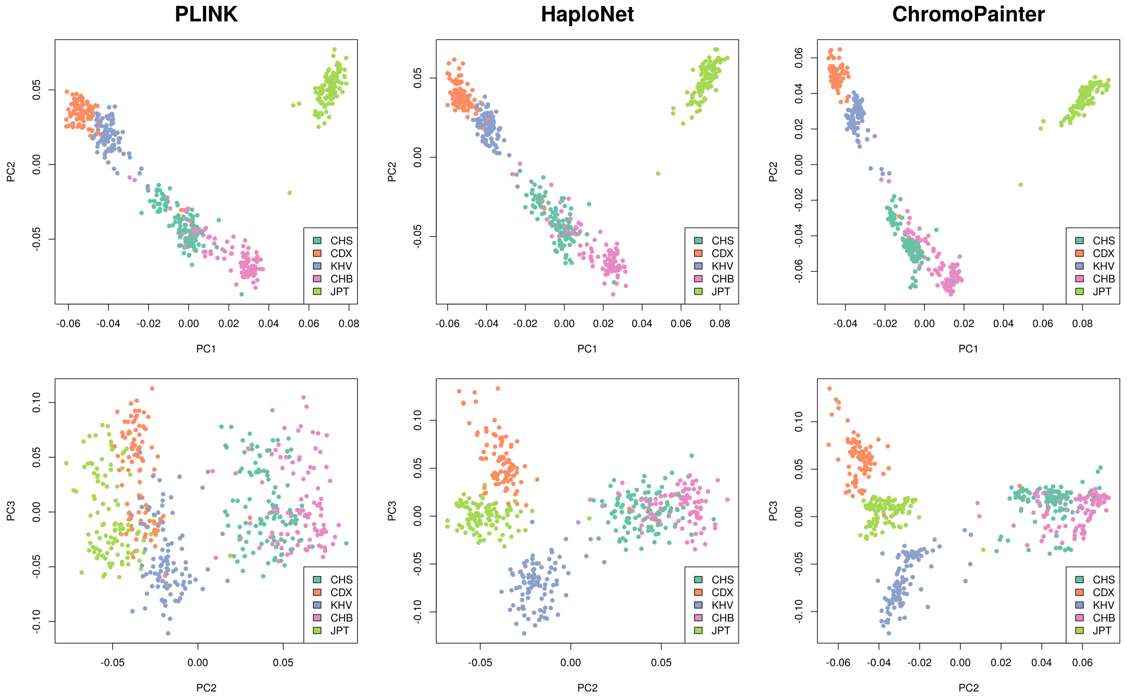


**Figure S12:** Inferred population structure in the EAS super population based on PCA using `PLINK` (left column), `HaploNet` (middle column), and `ChromoPainter` (right column). First row displays PC1 plotted against PC2, while the second row shows PC2 against PC3.

##### S7.2.5 EUR

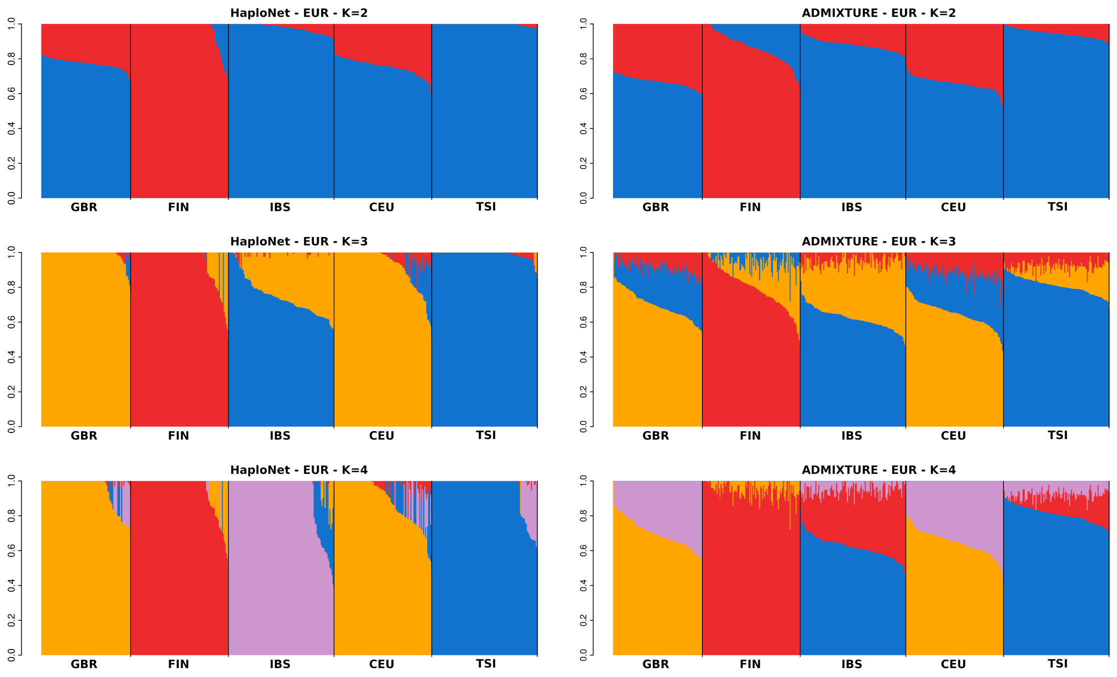


**Figure S13:** Estimated ancestry proportions in the EUR super population of the 1000 Genomes Project using `HaploNet` (left column) and `ADMIXTURE` (right column) for $K = 2, 3, 4$.

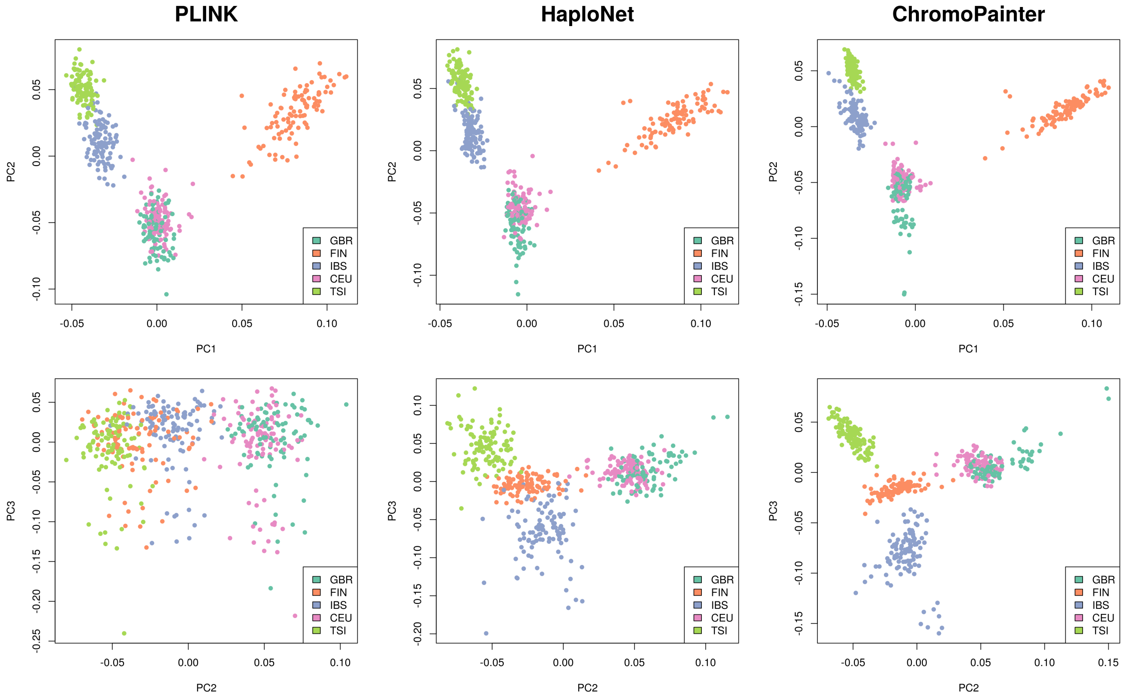


**Figure S14:** Inferred population structure in the EUR super population based on PCA using `PLINK` (left column), `HaploNet` (middle column), and `ChromoPainter` (right column). First row displays PC1 plotted against PC2, while the second row shows PC2 against PC3.

##### S7.2.6 SAS

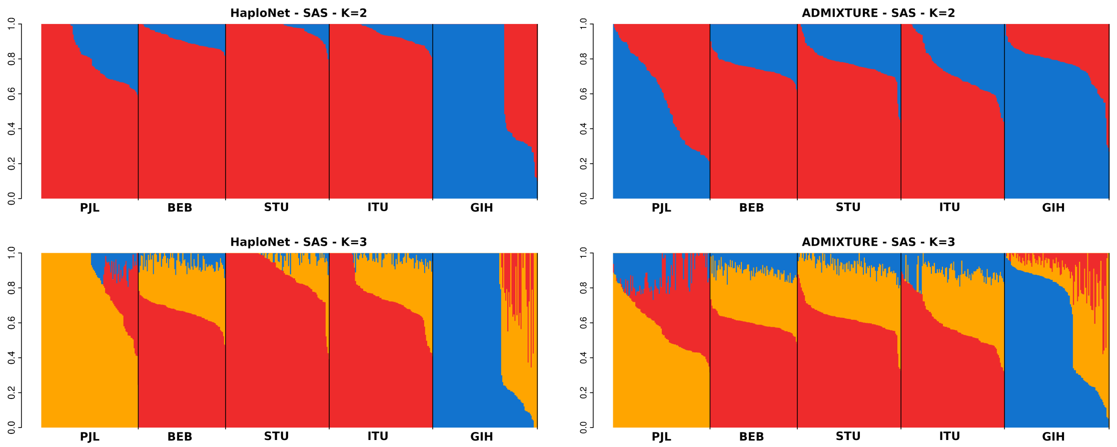


**Figure S15:** Estimated ancestry proportions in the SAS super population of the 1000 Genomes Project using `HaploNet` (left column) and `ADMIXTURE` (right column) for $K = 2, 3$.

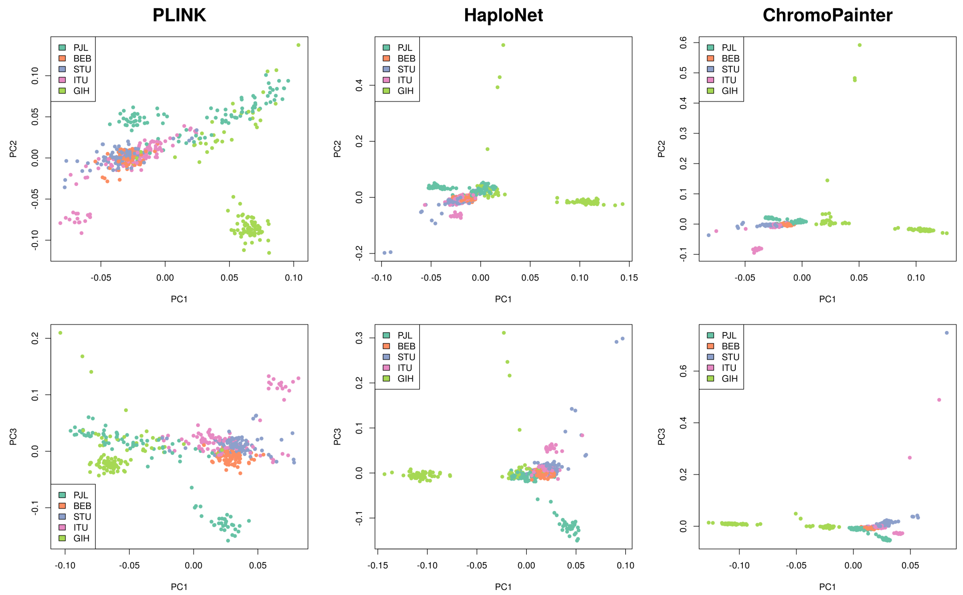


**Figure S16:** Inferred population structure in the SAS super population based on PCA using `PLINK` (left column), `HaploNet` (middle column), and `ChromoPainter` (right column). First row displays PC1 plotted against PC2, while the second row shows PC1 against PC3.

##### S7.2.7 Window size in HaploNet

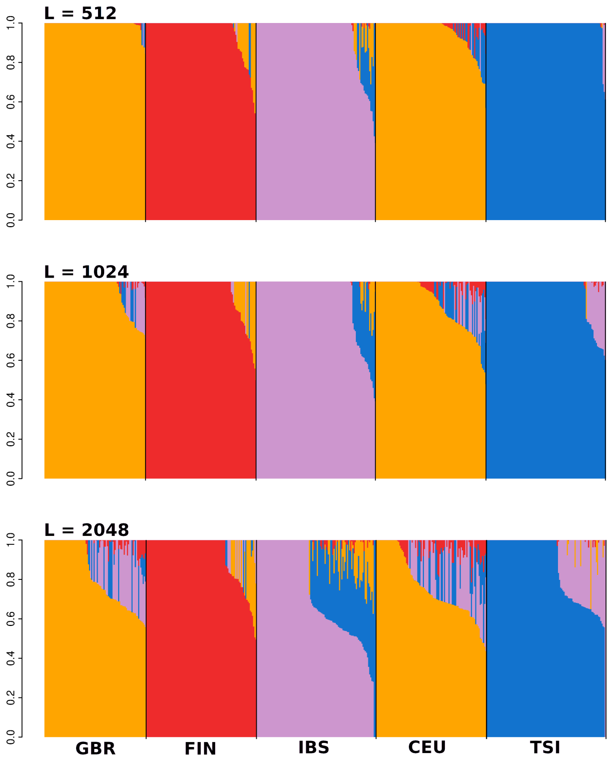


**Figure S17:** Estimated ancestry proportions in the EUR super population of the 1000 Genomes Project using `HaploNet` for $K = 4$ with three different window sizes ($L = 512, 1024, 2048$).

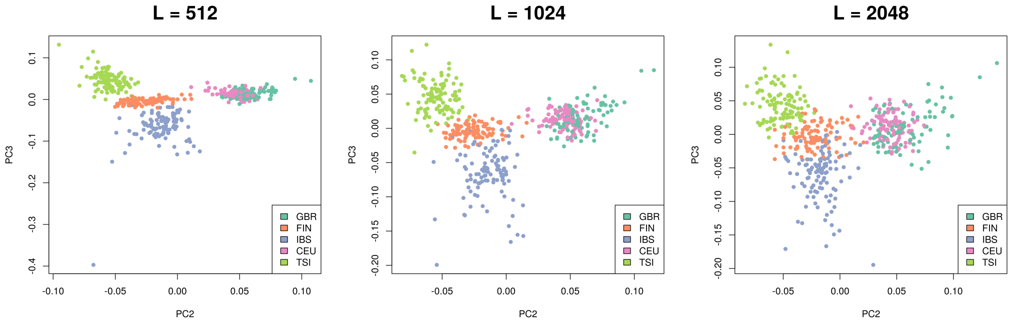


**Figure S18:** Inferred population structure in the EUR super population based on PCA using `HaploNet` with three different window sizes, $L = 512$ (first column), $L = 1024$ (second column) and $L = 2048$ (third column). The plots display PC2 plotted against PC3.

##### S7.2.8 Only categorical model in `HaploNet`

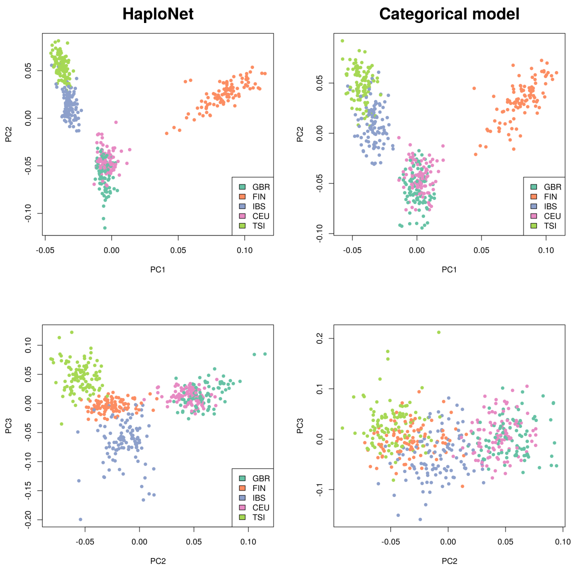


**Figure S19:** Inferred population structure in the EUR super population based on PCA using a simplified version of `HaploNet` that only performs haplotype clustering. The PCA plots of the full model (`HaploNet`) are shown in the left column while the PCA plots of the simplified model are shown in the right column.

##### S7.2.9 Permuted SNPs (no LD) for `HaploNet`

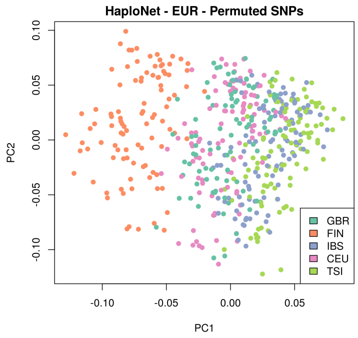


**Figure S20:** Inferred population structure in the EUR super population based on PCA using `HaploNet` on a randomly permuted haplotype matrix to break down LD structure. The plot displays PC2 plotted against PC3.

##### S7.2.10 EUR - Omni-chip sites

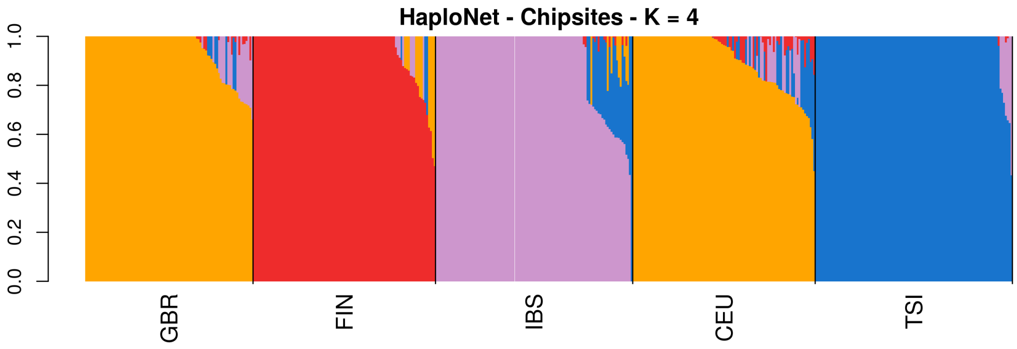


**Figure S21:** Estimated ancestry proportions in the EUR super population of the 1000 Genomes Project using `HaploNet` for $K = 4$ based on the high density genotype chip of the Omni platform used in the 1000 Genomes Project.

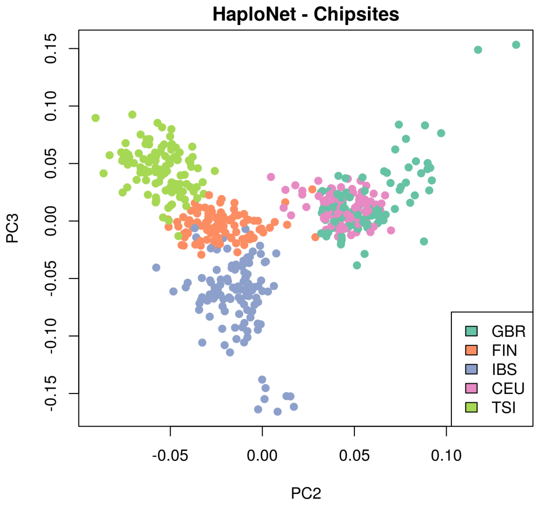


**Figure S22:** Inferred population structure in the EUR super population based on PCA using `HaploNet` based on the high density genotype chip of the Omni platform used in the 1000 Genomes Project. The plot displays PC1 plotted against PC2.

###### S7.2.11 Robustness

| | **Ancestry** | **PCA** |
|---|---|---|
| $L = 512$ | 14.47082 | 4.61830 |
| $L = 2048$ | 3.78338 | 3.20814 |
| Categorical model | - | 2.43939 |
| Permuted SNPs | - | 1.30697 |
| Omni-chip | 8.48543 | 3.75669 |
| $L = 1024$ (default) | 12.64427 | 3.93642 |

**Table S5:** Signal-to-noise ratio measures of various changes to `HaploNet` hyperparameters, model and input data for the EUR super population of the 1000 Genomes Project. The signal-to-noise ratio refers to mean between-population distance relative to the mean within-population distance of the estimated ancestry proportions and the principal components from PCA, respectively. Ancestry proportions have not been estimated for two scenarios due to their poor performance using PCA.

##### S7.2.12 Full dataset

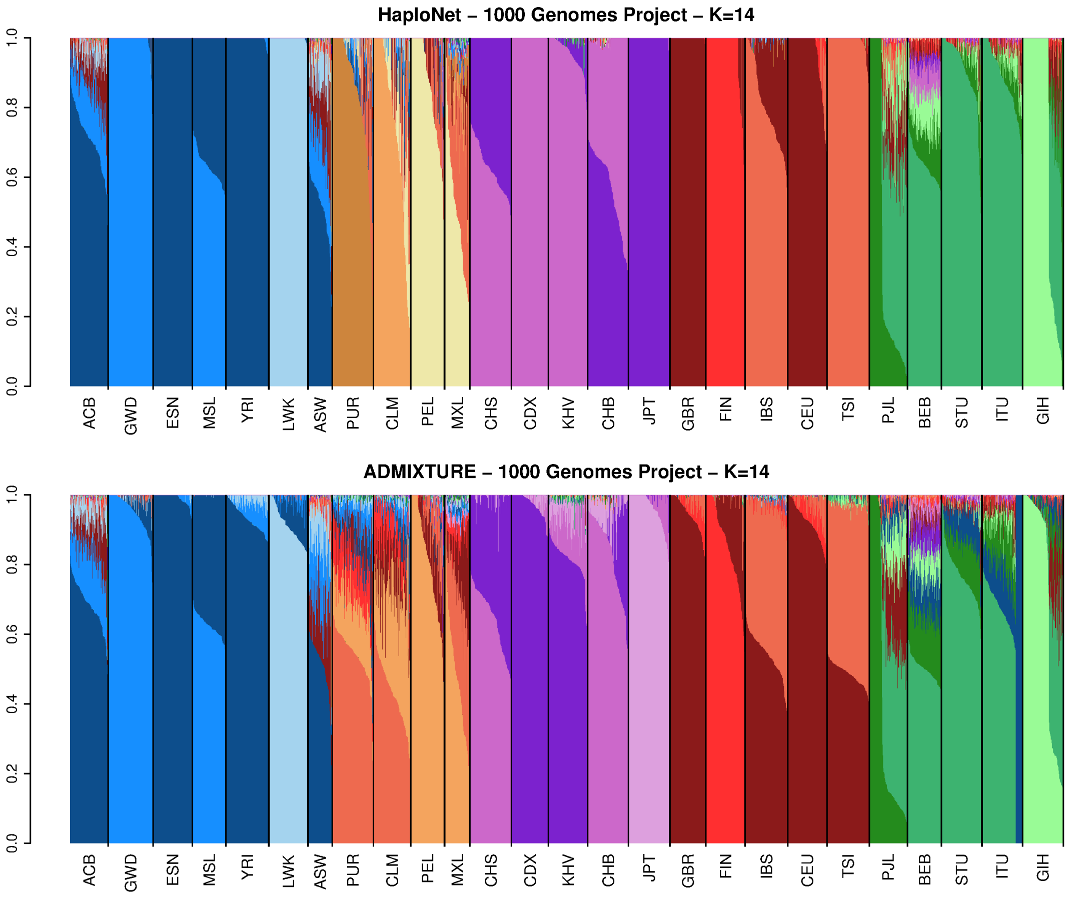


**Figure S23:** Estimated ancestry proportions in the full 1000 Genomes Project using `HaploNet` (top) and `ADMIXTURE` (bottom) for $K = 14$. `ADMIXTURE` has been performed on the LD pruned dataset for computational reasons and has reached convergence, however, it has had problems as seen especially for ITU.

#### S7.3 UK Biobank

##### S7.3.1 UK Biobank filtering

**Samples**

- Self-reported 'White-British' ethnicity (Data-field 21000)
- Similar genetic ancestry on basis of PCA (Data-field 22006)
- No kinship to other participants (Data-field 22021)

**Sites**

The sites filtering is mainly based on the SNP quality control process shared by the UK Biobank resource (file[2]).

- Sites on UK Biobank Axiom Array
- Sites passing all batch QC filters
- Removed sites in long range high LD regions (list)
- Sites with minor allele frequency $> 0.01$
- Sites with missingness rate $< 0.1$

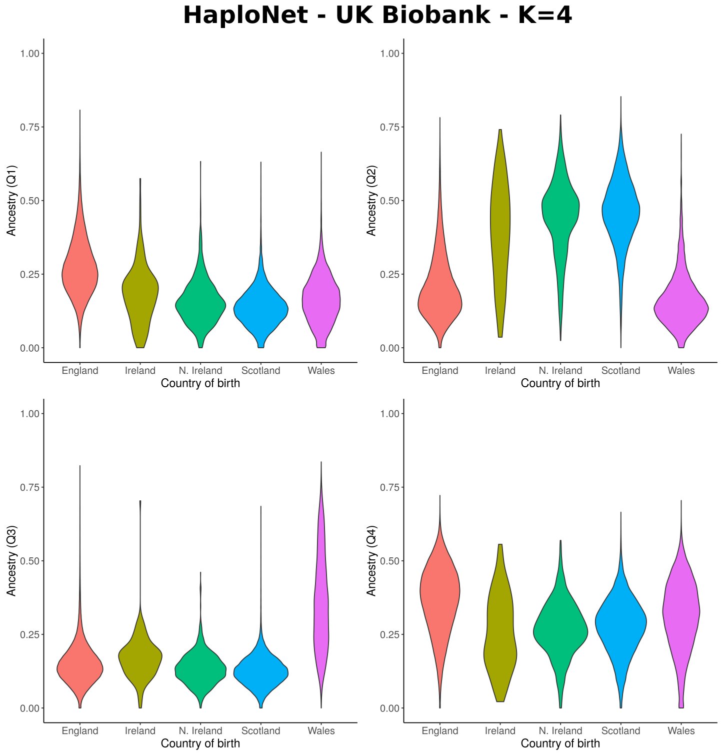


**Figure S24:** Ancestry proportions in the subset of unrelated self-identified 'White-British' using `HaploNet` for $K = 4$. Individuals have been grouped by their self-reported country of birth.

[2]The list of SNPs after strict LD pruning used for an analysis with PLINK is in field 'in_PCA'.

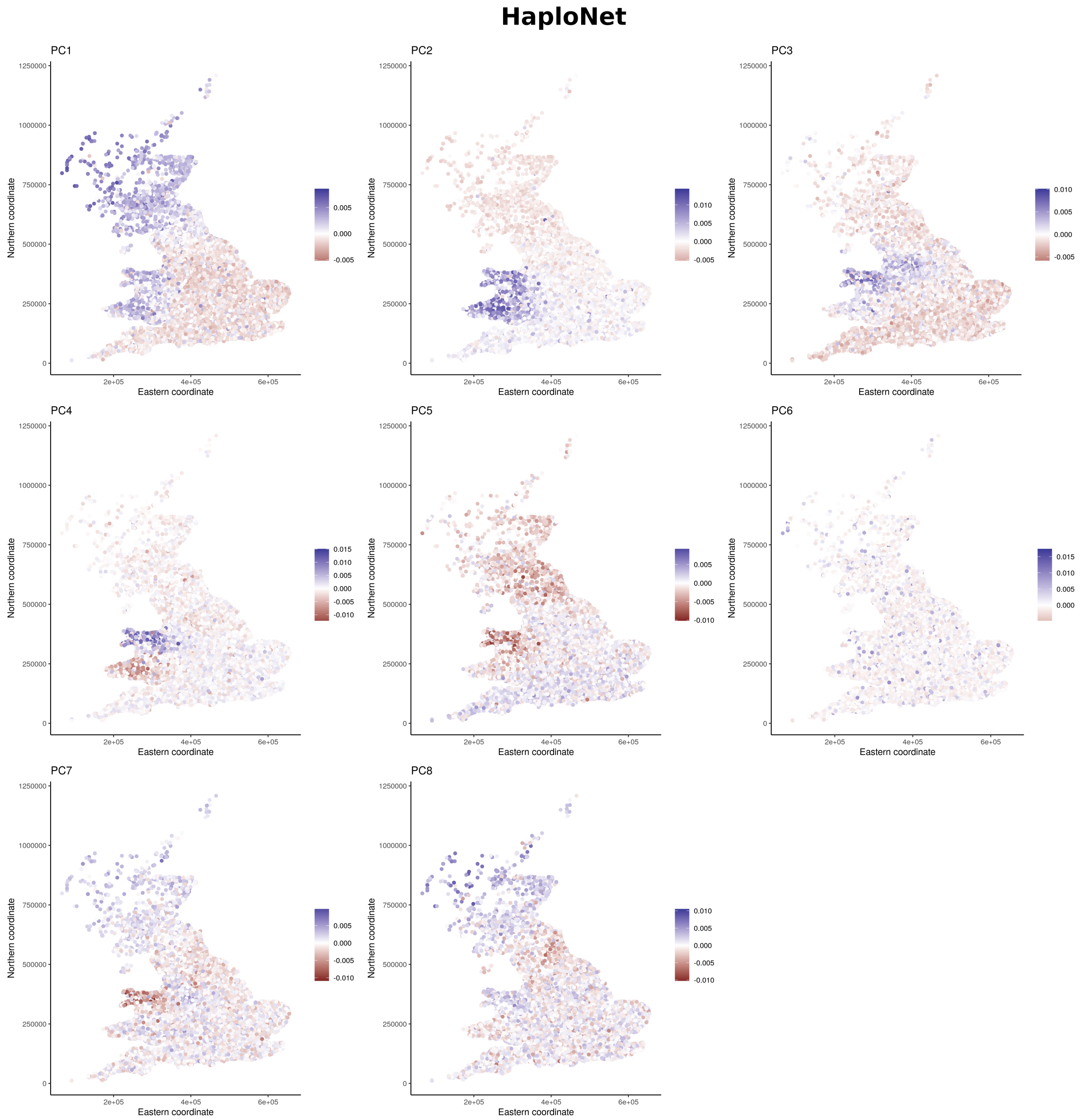


**Figure S25:** Inferred population structure in the subset of unrelated self-identified 'White-British' based on PCA using `HaploNet`. Individuals are plotted by their birthplace coordinates and colored by their inferred PC scores from PC1-PC8.

##### S7.3.2 Loadings

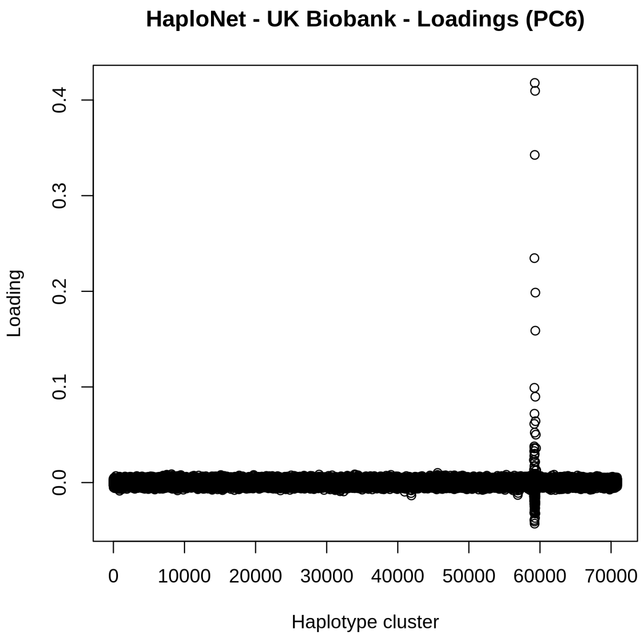


**Figure S26:** Inferred loadings of SNPs for PC6 using `HaploNet` on the UK Biobank dataset. The outliers span a region of 11.6 Mb on chromosome 16 (chr16:63991874-75563746).

#### References


[1] 1000 Genomes Project Consortium et al. A global reference for human genetic variation. *Nature*, 526(7571):68–74, 2015.

[2] D. H. Alexander, J. Novembre, and K. Lange. Fast model-based estimation of ancestry in unrelated individuals. *Genome research*, 19(9):1655–1664, 2009.

[3] C. Bycroft, C. Freeman, D. Petkova, G. Band, L. T. Elliott, K. Sharp, A. Motyer, D. Vukcevic, O. Delaneau, J. O'Connell, et al. The uk biobank resource with deep phenotyping and genomic data. *Nature*, 562(7726):203–209, 2018.

[4] E. Jang, S. Gu, and B. Poole. Categorical reparameterization with gumbel-softmax. *arXiv preprint arXiv:1611.01144*, 2016.

[5] J. Kelleher and K. Lohse. Coalescent simulation with msprime. *Statistical Population Genomics*, page 191, 2020.

[6] D. P. Kingma and M. Welling. Auto-encoding variational bayes. *arXiv preprint arXiv:1312.6114*, 2013.

[7] C. J. Maddison, A. Mnih, and Y. W. Teh. The concrete distribution: A continuous relaxation of discrete random variables. *arXiv preprint arXiv:1611.00712*, 2016.

[8] C. J. Maddison, D. Tarlow, and T. Minka. A* sampling. In *Advances in Neural Information Processing Systems*, pages 3086–3094, 2014.
